# Evidence for Stepwise Disruption of *E. coli* RNA Polymerase - λP_R_ Promoter Contacts and Bubble Collapse in Transcription Initiation

**DOI:** 10.1101/2024.11.22.624876

**Authors:** Max Rector, Renxi Li, Hao-Che Wang, M Thomas Record

## Abstract

In transcription initiation by *E. coli* RNA polymerase (RNAP), translocation of the RNA-DNA hybrid disrupts RNAP-promoter and σ^70^-core RNAP contacts, releasing σ^70^ and allowing RNAP to escape. Previously, to investigate whether RNAP-promoter contacts break step-by-step as the hybrid lengthens or concertedly at escape, we determined rate constants and activation energies of nucleotide-incorporation steps involving translocation at the λP_R_ promoter. Trends in these quantities with hybrid length were inconsistent with concerted models and provided evidence for stepwise disruption of RNAP-promoter contacts and collapse of the upstream bubble (−1 to −11). Here we report kinetic *m*-values quantifying urea and glycine betaine (GB) effects on rate constants of individual nucleotide-incorporation steps and compare with *m*-values predicted from structural and model compound information. GB is predicted to favor and urea disfavor binding the initiating NTPs and trigger-helix formation in 2-mer (pppApU) synthesis, largely because of burial of phosphate oxygens in NTP binding and amide oxygens in trigger-helix formation. Consistent with these predictions, GB accelerates and urea retards steps synthesizing 3-mer and 4-mer. However, both solutes retard mid-initiation steps (5-mer to 9-mer synthesis) where −10 contacts are proposed to break, exposing DNA phosphates and allowing the upstream initiation bubble to collapse. Strikingly, urea greatly accelerates while GB retards the 11-mer synthesis step, where −35-contacts are proposed to break, σ^70^ is released and RNAP escapes. Urea and GB kinetic *m*-values and activation energies of these steps are inconsistent with concerted models and support a stepwise model of contact disruption and bubble collapse in initiation.

## Introduction

Mechanistic studies with bacterial RNA polymerases (RNAP) using kinetic [e.g. 1–21] and structural [e.g. 22–33] approaches are revealing how the RNAP-promoter DNA complex operates as a molecular machine in open complex formation and subsequent nucleotide-incorporation steps of transcription initiation. Kinetic-mechanistic studies of open complex formation and initiation by bacteriophage T7 RNAP [e.g. 34–39] and mitochondrial RNAP [40] and of elongation by bacterial [e.g. 41–44], T7 phage [e.g. 45] and eukaryotic [e.g. 46–49] RNAP have also been performed to complement the structural information about mechanism [e.g. 50–61] obtained for those systems. While substantial progress has been made, much remains to be learned about the steps of bacterial transcription initiation and the similarities and differences in initiation mechanisms between bacterial and other RNAP.

Our focus is on major conformational changes and disruption of interfaces in the *E. coli* σ^70^ RNAP-λP_R_ promoter initiation complex that occur as a result of translocation of the DNA template relative to the active site. These changes result in escape of RNAP from the promoter and the transition to elongation. One downstream base pair is opened in each translocation step. Translocation scrunches the upstream bubble strands [20, 21] and causes unfavorable steric interactions between the growing RNA-DNA hybrid and RNAP [29], stressing upstream contacts. The free energy costs of downstream base pair opening and upstream stress make translocation unfavorable in each step of initiation. The incoming NTP binds only to the post-translocated state, making that state more favorable [42, 62–64]. RNAP interactions with the strands of the discriminator and −10 regions and with the upstream duplex are disrupted as a result of translocation stress, allowing bubble collapse and duplex formation by the strands of the upstream initiation bubble (−1 to −11 for λP_R_). At the λP_R_ promoter, the σ^70^ specificity subunit is displaced and core RNAP escapes from the promoter during the step synthesizing an ∼11-mer RNA [17, 65], marking the end of the initiation phase of transcription.

Not all initiating complexes in the population are able to progress to the escape point and make the transition to elongation, however. A substantial population (30-50%) of initiating complexes, called nonproductive, stall before the escape point after synthesis of a short RNA (< 11-mer at the λP_R_ promoter), presumably because of difficulty in translocation [12-14, 17, 66, 67]. In some cases (e.g. higher temperatures, shorter RNA), the nascent RNA dissociates from the stalled nonproductive complex but RNAP does not, returning to the TSS and reinitiating in a cycle of abortive initiation.

Here we address the questions of whether disruption of RNAP-promoter contacts and bubble collapse occur together with σ^70^ release in the RNAP escape step (the concerted model), or whether these changes in conformation and interfaces occur in both mid- and late-initiation steps (the stepwise model). Previous structural, biochemical, and kinetic-mechanistic research on initiation by the phage T7 RNAP revealed that contacts with three promoter regions (the bubble template strand, the specificity and AT-rich duplex regions) are disrupted sequentially in multiple nucleotide-incorporation steps as the RNA-DNA hybrid extends to ∼12 bp, ending in RNAP escape [38, 39, 51-54; see Discussion]. Kinetic-mechanistic studies of initiation by *E. coli* RNAP at the λP_R_ promoter led to the proposal that contacts with three promoter regions (discriminator strands, −10 strands and upstream (extended −10, −35) duplex) are also disrupted sequentially in multiple steps as the RNA-DNA hybrid extends to ∼11 bp [13, 14]. Here we test this proposal by determining the effects of two solutes on the kinetics of individual initiation steps.

To probe whether interactions between RNAP and promoter DNA in a productive initiation complex are disrupted step-by-step or concertedly, we quantify the effects of urea and glycine betaine (GB) on the kinetics of individual steps. These solutes were chosen because their interactions with the different types of protein and nucleic acid surface are quite well-characterized and this information has proved useful to interpret and/or predict their often-large effects on protein and nucleic acid processes. Urea, GB and/or other solutes were previously used in studies of the kinetics and mechanisms of forming the RNAP-promoter initiation complex [4, 68], as well as protein folding [69] and formation of a lac repression complex [70]. In these previous studies, solute effects on the kinetics were interpreted in terms of solute interactions with the biopolymer surface that is exposed or buried in the interactions and conformational changes that convert reactants to the highest free energy transition state. New information was obtained about these transition states and the mechanisms of these processes, not readily available by other methods. A summary of the analysis and interpretation of solute kinetic *m*-values in terms of changes in water-accessible surface area (i.e ΔASA) is provided in SI.

## Results

### Urea and GB Effects on Rates of Steps of RNA-DNA Hybrid Extension in Transcription Initiation: Rapid Quench-Flow and Gel Separation Assays

Initiation kinetics experiments were performed as a function of urea and GB concentration by mixing preformed RNAP-λP_R_ open complexes (OC) with initiation solution containing ATP, UTP, GTP, heparin, and either urea or GB (see SI Methods) at 19 °C in a rapid quench-flow (RQF) mixer. Conditions and most details of these experiments were the same as those used previously to study initiation kinetics in the absence of urea and GB [12–14, 17], except as noted in SI Methods. In all cases, the promoter template strand (ending at +41 relative to the +1 transcription start site) was modified from that of wild-type λP_R_ so that CTP is not needed to synthesize a 16-mer RNA. The first requirement for CTP is at position +17, and subsequently at +32. Heparin is present in the initiation solution to bind free RNAP and prevent re-initiation.

**Figure 1** shows representative polyacrylamide gel separations of RNA oligomers present as a function of time (from 0.1 s to 120 s) in initiation experiments performed at 200 μM ATP and UTP (the two initiating NTP) and 10 μM GTP (the α-^32^P-labeled NTP). **Panel A** is a control experiment at this NTP condition in the absence of added solute. **Panels B** and **C** are at final solute concentrations of 0.5 M urea and 0.5 M GB, respectively. Examples of initiation gels at 1 M urea and GB are shown in **SI Figure S1**.

These panels of **Figure 1** reveal the kinetics of stepwise growth of the RNA-DNA hybrid beginning with synthesis of 3-mer (pppApUpG). No information is obtained about the initial dinucleotide pppApU, which is not labeled in experiments with α-^32^P-labeled GTP and is not present in sufficient amounts to be detected in experiments using α-^32^P-labeled UTP. Any labeled pppApU in the latter assays apparently co-migrates with a labeled contaminant band in the UTP stock for the gel conditions used.

**Figure 1.**
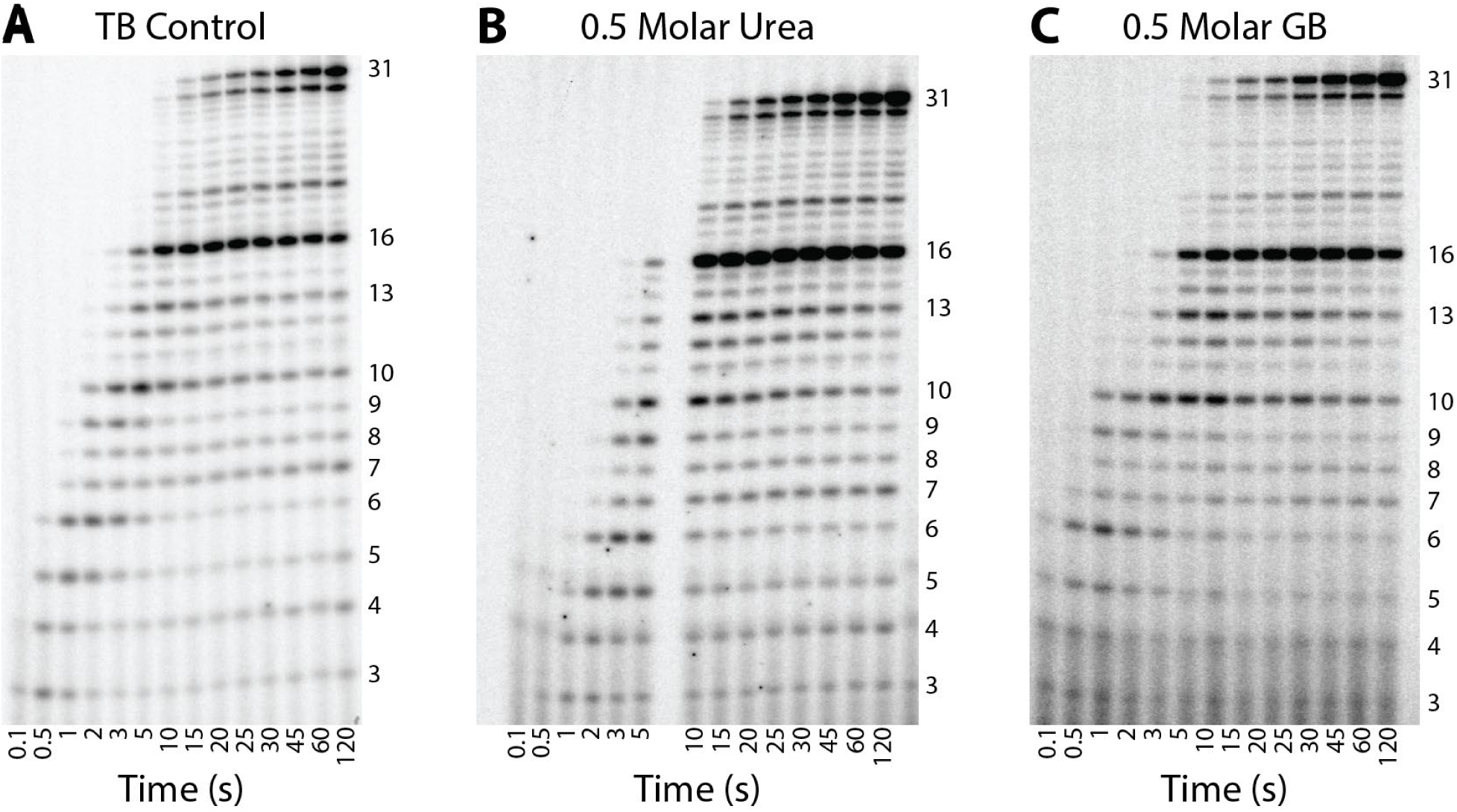
Time courses of transcription initiation by E. coli RNA polymerase-λP_R_ promoter complexes at 0.5 M urea and 0.5 M GB. Representative gel separations of short (pre-escape; <11-mer)) and long (≥11-mer) RNA transcripts as a function of time (0.1 s to 120 s) after NTP addition at the high UTP condition at 19 °C A) Control (TB, no added solute); B) 0.5 M urea; C) 0.5 M glycine betaine (GB).

**Figures 1** and **S1** reveal the existence of two classes of initiating complexes (productive and nonproductive) at all solute conditions investigated and demonstrate two kinetic phases of RNA synthesis by the population of initiating complexes. Both of these observations are completely consistent with those made previously in the absence of added solute [12–14, 17].

Productive complexes synthesize a 16-mer RNA relatively rapidly (∼10 s for the conditions of Figs 1 and S1). In that first 10 s, amounts of short RNAs increase and then decrease as the next larger RNA is synthesized, while the amount of RNA ≥11-mer increases monotonically. Gels exhibit slow readthrough of the CTP-requiring stop at +17, as observed previously, resulting in synthesis of longer RNA oligomers up to the second CTP stop (31-mer synthesis). No readthrough of this second C-stop is observed, presumably because the transcription rate is greatly reduced near the end of the template fragment. RNAP in productive complexes escapes from its contacts with the λP_R_ promoter in synthesizing 11-mer RNA from 10-mer [17, 65]. All RNA lengths ≥11-mer are considered together as “full-length” (post-escape) RNA in our analysis, so readthrough has no effect on the analysis.

As previously reported [12-14, 17, 66, 67], short RNAs observed in these gels at times greater than 10 s are primarily those synthesized by nonproductive complexes before stalling. Slow increases in some short RNA populations with increasing time beyond 10 s are from stalled nonproductive complexes that release their RNA and reinitiate (abortive initiation). At all solute concentrations investigated, about half of λP_R_ open complexes are productive, as observed previously in the absence of added solutes [12–14, 17].

Because there are multiple sites for incorporation of an α-^32^P-labeled nucleotide (in place of the corresponding non-radioactive nucleotide) in all RNA oligomers longer than 5-mer (when labeling with α-^32^P-GTP) or 3-mer (when labeling with α-^32^P-UTP), gels like those in **Figure 1** give a skewed picture of the relative mole amounts of different RNA species present. Band intensities of each RNA oligomer are therefore corrected using incorporation probabilities [14] to obtain mole amounts, which are then normalized to the total mole amount of ≥11-mer RNA synthesized at long times in that experiment (see SI Methods). Averages of mole amounts of each oligomer length are plotted vs time in **Figure 2** for experiments without added solute (**panel A**) and at 0.5 M urea (**panel B**) and GB (**panel C**). **Figure S2** shows the corresponding behavior in experiments at 1 M urea (**panel B**) and GB (**panel C**).

**Figure 2.**
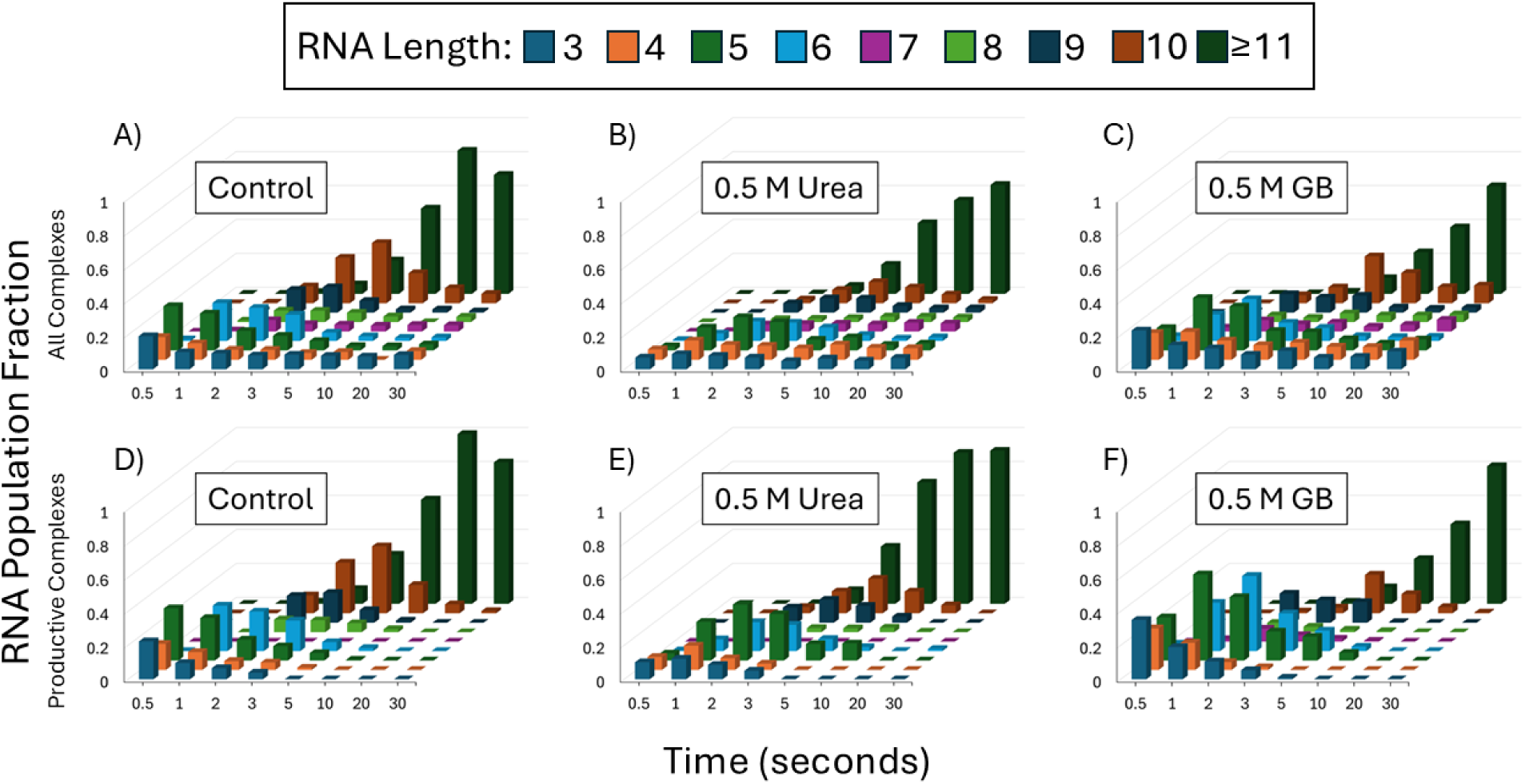
Time-evolution of populations of short RNAs (pre-escape; 3-mer to 10-mer) and of long (≥11-mer) RNA) in initiation at λP_R_ promoter at 0.5 M Urea and 0.5 M GB. NTP condition: high ATP, UTP, low GTP**. Panels A-C:** Total RNA from both productive and non-productive complexes. **Panels D-F**: RNA from productive complexes only. **Panels A, D:** Control (TB, no added solute). **Panels B, E:** 0.5 M urea. **Panels C, F:** 0.5 M glycine betaine. For all panels, all RNA amounts are normalized to the final amount of long (≥11-mer) RNA.

**Figures 2** and **S2** reveal that shorter intermediates (3-mer to 8-mer) in RNA synthesis by productive complexes dominate the kinetics for the first 3-5 s, both with and without added solute. In urea the synthesis of these shorter intermediates occurs somewhat more slowly than for the no-solute control or in GB. Longer intermediates (9-mer, 10-mer) and full-length RNA synthesis dominate the later kinetics (3-30 s). Synthesis of these longer intermediates from shorter intermediates occurs more slowly in GB than in urea or the control.

## Analysis and Discussion

### Time Course of Transient Short RNA Intermediates in Long RNA Synthesis By Productive RNAP-Promoter Complexes

Results like those in **Figures 1** and **2 (Panels A-C),** as well as **Figures S1** and **S2 (Panels A-C),** for amounts of each RNA length vs. time in synthesis by both productive and nonproductive complexes were analyzed as described previously to remove contributions from short RNA synthesis by nonproductive complexes [12-14; see SI Methods]. The resulting time courses for productive complexes are shown in the bar graphs of **Figure 2** for 0 and 0.5 M urea (**panels D, E**) and GB (**panel F**) and in **Figure S2 panels D-F** for 0 and 1 M urea and GB. These clearly reveal the progressive build-up and decay of each intermediate RNA length as well as the initial lag and subsequent buildup of long (≥11-mer) RNA. Compared with the 0 M control, urea (**Figure 2 Panel E, Figure S2 Panel C**) shifts the transient peaks for the shortest RNAs to longer times but has less effect on the rate of long RNA synthesis. GB (**Figure 2 Panel F, Figure S2 Panel D**), on the other hand, has little effect on the timing of the transients for the shortest RNAs, but slows the synthesis of long RNA. Therefore, urea retards early steps and accelerates later steps of initiation, while GB has the opposite effects.

### Overall 2^nd^ Order Rate Constants for Initiation Steps as Functions of Urea and GB Concentration

Translocation in initiation at the λP_R_ promoter is intrinsically unfavorable and reversible [13, 14]. The incoming NTP binds only to the post-translocated state [42, 62–64], thereby increasing the population fraction of post-translocated complexes. This coupling of translocation and NTP binding makes NTP binding weaker and increases the apparent NTP Michaelis constant K_m_ for initiation steps. Consequently overall 2^nd^ order rate constants for each step of nucleotide incorporation into RNA involving translocation (designated k_obs,i_ for step i and analogous to overall 2^nd^ order rate constants k_cat_/K_m_ in traditional enzyme kinetic analyses) are sufficient to characterize the kinetics at the NTP concentrations investigated [12–14].

As a first level of analysis, kinetic data at each solute concentration including those in **Figures 1-2** and **Figures S1-S2** were fit to the minimal 11-step mechanism of initiation, previously shown to be sufficient for the λP_R_ promoter at 19 °C [13, 14]. Values of k_obs,i_ are obtained for each step of nucleotide incorporation involving translocation (steps 3 to 11). In these fittings, as in previous analyses, rate constants for the steps of pppApU (2-mer) synthesis were fixed. As described in SI Methods, rate constants for binding the two initial NTP and for TL folding in 2-mer synthesis at each urea and GB concentration were obtained from those used previously in the absence of these solutes using *m*-values predicted from structural analysis and model compound information. Urea is predicted to disfavor NTP binding and TL folding in 2-mer synthesis; GB is predicted to favor these processes. Fixing the rate constants for steps of 2-mer synthesis, instead of floating these quantities, reduces the uncertainties in fitted rate constants k_obs,i_ for subsequent steps. Values of k_obs,i_ for these subsequent steps are relatively insensitive to the choices of urea and GB *m*-values for 2-mer synthesis, as detailed in SI Methods.

Semi-log plots of rate constants k_obs,i_ (obtained as above from non-global fitting) as a function of urea and GB concentration for incorporation of the 3^rd^ (G), 7^th^ (U), and 11^th^ (G) nucleotide, representative of early, mid- and late initiation steps, are shown in the panels of **Figure 3**. These plots are linear, as is generally observed for solute effects, indicating from **Eq. S6** that urea and GB kinetic *m*-values for these steps do not depend significantly on solute concentration. More accurate kinetic *m*-values of these and other initiation steps are obtained by global fitting of these data, as described below.

**Figure 3.**
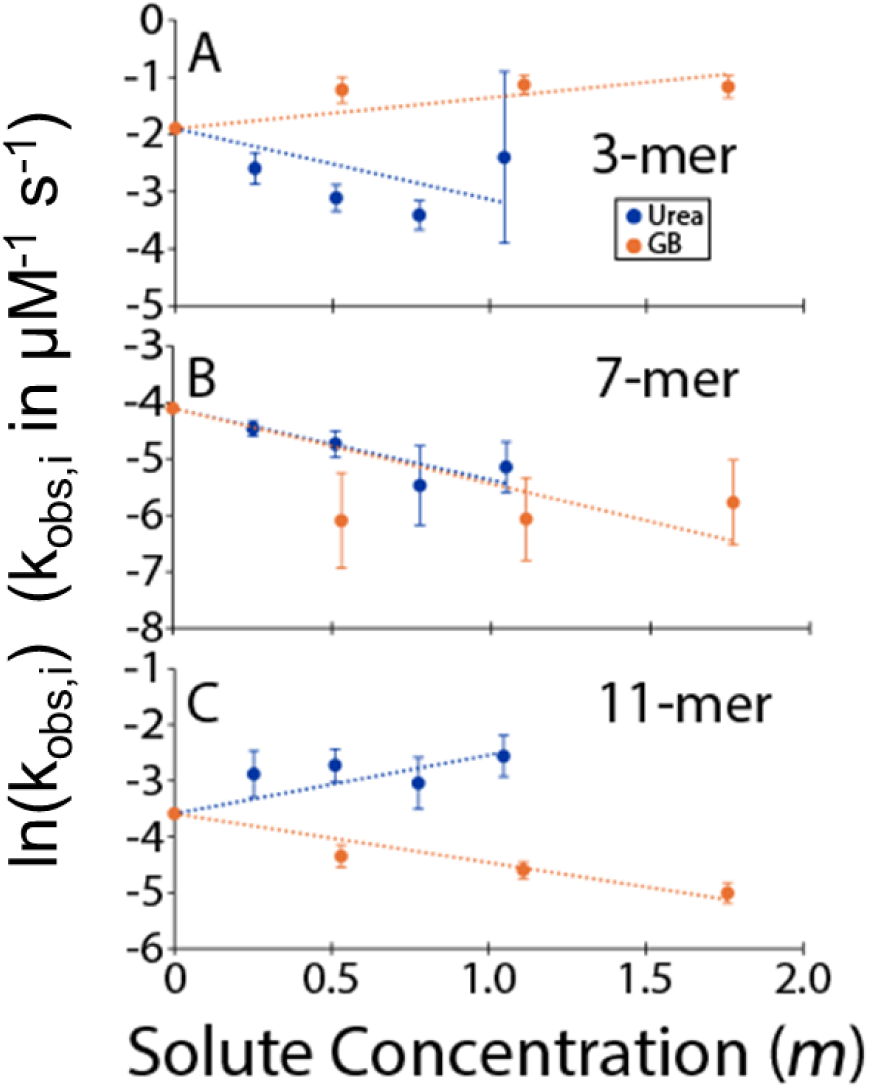
Behavior of 2^nd^ Order Nucleotide Incorporation Rate Constants (k_obs,i_) as Functions of Urea and GB Concentration. Natural logarithms of rate constants k_obs,i_, obtained from non-global fitting as described in SI Methods, plotted as a function of concentration of urea (blue) or GB (orange). Linear fits (with slopes equal to – kinetic *m*-value/RT) to urea and GB ln(k_obs,i_) values in each panel are constrained to have a common intercept at zero solute concentration.

**Figure 3A** shows that GB increases the rate constant for incorporation of the 3^rd^ (G) nucleotide while urea reduces it. The observed urea *m*-value for this 3^rd^ step has the same sign and a similar magnitude to that predicted from structural information for incorporation of the 2^nd^ (U) nucleotide, while the GB *m*-value for incorporation of the 3^rd^ (G) nucleotide agrees in sign but is appreciably smaller in magnitude than the structural prediction for the 2^nd^ step. Strikingly, both solutes reduce the rate constant for incorporation of the 7^th^ (U) nucleotide (**Figure 3B**). Urea increases and GB reduces the rate constant for incorporation of the 11^th^ (G) nucleotide (**Figure 3C**). For many protein processes, urea and GB m-values are of opposite sign but similar magnitude. This behavior is observed for step 3 and, with a change of sign, step 11 but not for step 7, indicating that there are very significant differences in the amounts and types of RNAP and/or promoter surface area exposed or buried in these steps. Together, these results are qualitatively consistent with the observation from **Figure 2** that urea retards early steps of initiation but accelerates late steps, while GB has the opposite effect.

### Solute *m*-Values of Initiation Steps Obtained by Global Fitting

Global fitting of all kinetic data for each solute was performed to obtain kinetic *m*-values directly for each initiation step beyond pppApU synthesis. This fitting introduces the exponentiated version of **Eq. S6** relating rate constants of a step of nucleotide incorporation as a function of solute concentration to its kinetic *m*-value, with intercept terms determined by published values of rate constants in the absence of added solute [13, 14]. For the high UTP condition, time courses (log time scale) predicted by global fitting for 3-mer, 5-mer, 7-mer and 9-mer transient intermediates and for long (≥11-mer) RNA synthesis are plotted together with experimental results in **Figure 4 Panels A-C** for 0 and 0.5 M urea and 0.5 M GB. and compared with experimental results. **Figure S3 Panels A–C** show corresponding time courses for 0 and 1 M urea and 1 M GB.

**Figure 4.**
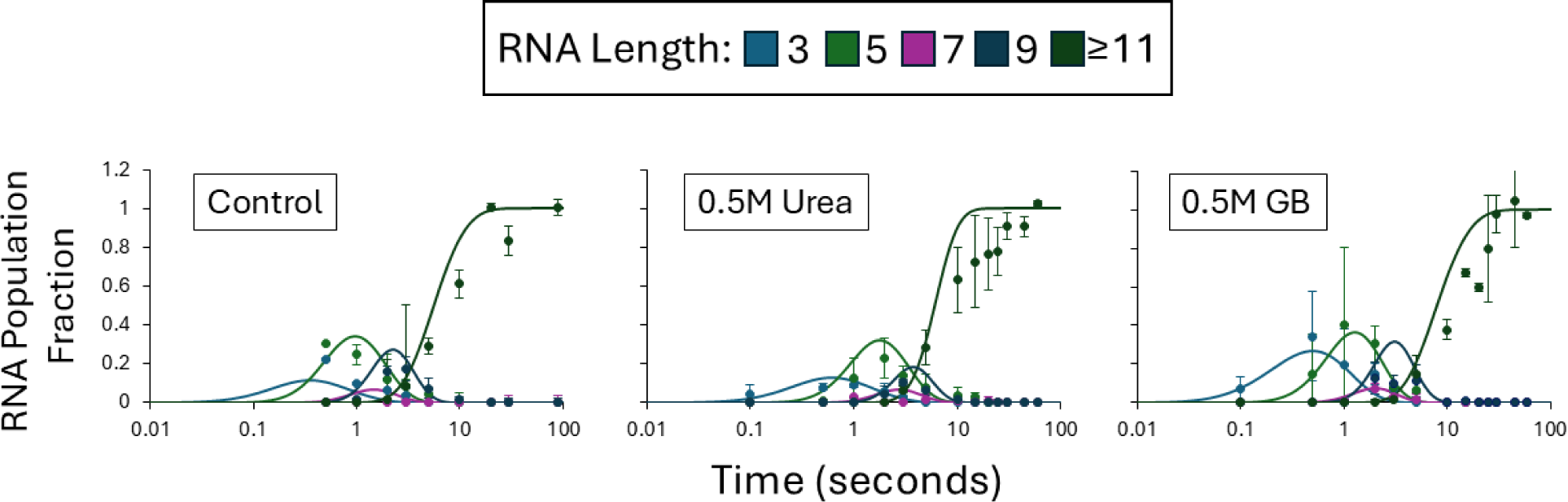
Fitting of time course data to the step-by-step initiation mechanism. Results for four pre-escape intermediates and for ≥11-mer RNA at 0.5 M urea, 0.5 M GB and the control (see **Figure 2** Panels D-F) are compared with curves predicted from a global fit on this log time plot. **Panel A):** No added solute. **Panel B:** 0.5 M urea. **Panel C:** 0.5 M glycine betaine. All RNA amounts are normalized as in **Figure 2**.

Urea and GB kinetic *m*-values of initiation steps 3-11 determined by global fitting are plotted with uncertainties in **Figure 5** and listed in **Table 1**. All these steps begin with reversible translocation, followed by reversible NTP binding and the rate determining step(s) of incorporation of the nucleotide into the RNA (TL folding, catalysis). In the following sections, we interpret these results to extract the contribution of translocation to the kinetic *m*-value of each step and obtain information about the translocation-linked processes of contact disruption and duplex formation that occur prior to and/or in escape of RNAP from the promoter.

**Figure 5.**
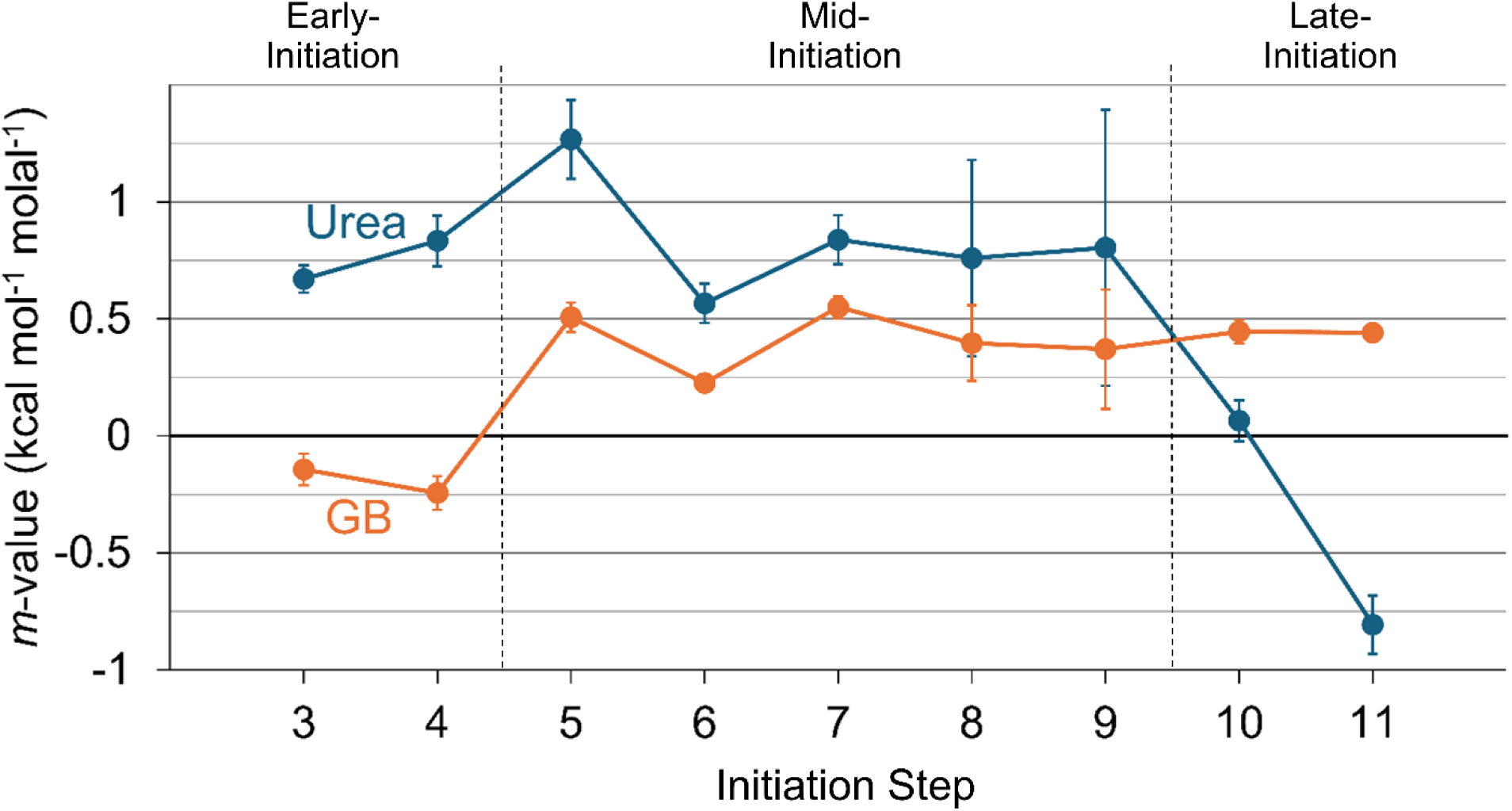
Urea and GB Kinetic m-Values for Early-, Mid- and Late-Initiation Steps at λP_R_ Promoter. Solute dependences (kinetic *m*-values, in kcal mol^-1^ molal^-1^) of 2^nd^ order nucleotide addition rate constants k_obs,i_ for urea (blue) and glycine betaine (orange) from **Table 1**, are plotted vs. initiation step (i.e. RNA-DNA hybrid length). These k_obs,i_ values were obtained by global fitting of initiation kinetic data for productive complexes as described in SI Methods.

**Table 1.** Urea and GB *m*-Values, Activation Energies and *Excess* Quantities for Initiation Steps with Translocation.

| Observed for step synthesizing | Urea $m$-value <sup>a</sup> | Excess Urea $m$-value <sup>a,b</sup> | GB $m$-value <sup>a</sup> | Excess GB $m$-value <sup>a,c</sup> | $E_{act}^d$ | Excess $E_{act}^d$ |
| --- | --- | --- | --- | --- | --- | --- |
| 3-mer | $0.67 \pm 0.06$ | - | $-0.17 \pm 0.07$ | - | $27 \pm 14$ | - |
| 4-mer | $0.83 \pm 0.11$ | - | $-0.31 \pm 0.08$ | - | $26 \pm 2$ | - |
| 5-mer | $1.27 \pm 0.17$ | $0.5 \pm 0.2$ | $0.52 \pm 0.06$ | $0.7 \pm 0.1$ | $5.1 \pm 10$ | $-21 \pm 12$ |
| 6-mer | $0.57 \pm 0.08$ | $-0.2 \pm 0.2$ | $0.25 \pm 0.04$ | $0.5 \pm 0.1$ | $7.2 \pm 3$ | $-19 \pm 7$ |
| 7-mer | $0.84 \pm 0.11$ | $0.1 \pm 0.2$ | $0.57 \pm 0.05$ | $0.8 \pm 0.2$ | $6.8 \pm 2$ | $-19 \pm 6$ |
| 8-mer | $0.76 \pm 0.42$ | $0.0 \pm 0.4$ | $0.42 \pm 0.15$ | $0.6 \pm 0.2$ | $6.6 \pm 7$ | $-20 \pm 9$ |
| 9-mer | $0.80 \pm 0.59$ | $0.0 \pm 0.6$ | $0.41 \pm 0.23$ | $0.6 \pm 0.3$ | $-4.4 \pm 10$ | $-30 \pm 12$ |
| 10-mer | $0.06 \pm 0.09$ | $-0.7 \pm 0.1$ | $0.47 \pm 0.05$ | $0.6 \pm 0.1$ | $16 \pm 4$ | $-10 \pm 7$ |
| 11-mer | $-0.81 \pm 0.12$ | $-1.6 \pm 0.2$ | $0.44 \pm 0.02$ | $0.6 \pm 0.1$ | $21 \pm 2$ | $-5 \pm 6$ |
| Sum (5 to 9-mer) | | $0.4 \pm 0.8$ | | $3.2 \pm 0.4$ | | $-109 \pm 21$ |
| Sum (10 to 11-mer) | | $-2.3 \pm 0.2$ | | $1.2 \pm 0.1$ | | $-15 \pm 9$ |
| Sum (5 to 11-mer) | | $-1.9 \pm 0.8$ | | $4.4 \pm 0.4$ | | $-124 \pm 23$ |

<sup>a</sup> kcal mol<sup>-1</sup> molal<sup>-1</sup>
<sup>b</sup> 3-mer/4-mer average urea $m$ -value $0.75 \pm 0.12$ kcal mol<sup>-1</sup> molal<sup>-1</sup>;
<sup>c</sup> 3-mer/4-mer average GB $m$ -value $-0.24 \pm 0.11$ kcal mol<sup>-1</sup> molal<sup>-1</sup>.
<sup>d</sup> kcal mol<sup>-1</sup>. 3-mer/4-mer average activation energy $E_{act} = 26 \pm 6$ kcal mol<sup>-1</sup> (uncertainty assigned by averaging those for all positions)

### Step-by-Step Solute Kinetic *m*-Values Divide into Three Groups

Qualitatively **Figure 5** indicates that globally-fitted urea and GB kinetic *m*-values of initiation steps with translocation, considered together, divide into three groups: i) 3- and 4-mer synthesis, ii) 5-mer to 9-mer synthesis, and iii) 10-mer and 11-mer synthesis. We call these early-, mid- and late-initiation steps respectively, as shown in **Figure 5**. Results from non-global fitting for 3-mer, 7-mer and 11-mer in **Figure 3** match these three groupings. **Figure 5** also shows that previously-reported activation energies (E_act_), determined from the temperature dependences of rate constants of these steps, divide into three similar groups [13].

Urea kinetic m-values for early-initiation steps in **Figure 5** are unfavorable (positive, retarding) and of similar moderate magnitude, while GB kinetic *m*-values of these steps are favorable (negative, accelerating) and of similar moderate magnitude. These 3-mer and 4-mer kinetic *m*-values contain contributions from solute effects on opening a downstream base pair, scrunching the upstream strands, and on conformational changes in RNAP in translocation, NTP binding, trigger loop folding and other events of catalysis leading to the high free energy transition state. These contributions to kinetic *m*-values of steps 3 and 4 also occur in all subsequent steps of initiation. E_act_ values for these two steps were also found to be very similar (each approximately 26 kcal mol^-1^) [13]. These E_act_ values include contributions from downstream bp opening and NTP binding, as well as TL folding and catalysis, all of which also contribute to E_act_ values of subsequent steps. From the analysis of E_act_ values, we previously concluded that translocation in 3-mer and 4-mer synthesis did not involve any major (large enthalpy) coupled conformational changes [13].

Mid-initiation steps (steps 5-9) define a second region of solute kinetic *m*-values in **Figure 5**. GB kinetic *m*-values change from moderately favorable in steps 3 and 4 to moderately unfavorable in steps 5-9, while urea m-values change only modestly if at all from those of the earlier initiation steps. E_act_ values for these mid-initiation steps were found to be 20 to 30 kcal mol^-1^ smaller than those of steps 3 and 4, indicating that an additional enthalpically-favorable process occurs in mid-initiation steps [13].

Late-initiation steps 10 and 11, in which the σ^70^ subunit is released from the λP_R_ initiation complex and RNAP escapes from the λP_R_ promoter [17, 62], constitute a third region of **Figure 5**. Urea *m*-values shift dramatically from unfavorable for steps 3-9 to neutral for step 10 and favorable for step 11, while GB *m*-values for these two late steps are similarly unfavorable to those of mid-initiation steps. E_act_ values for steps 10 and 11 were found to greatly exceed those for steps 5-9, with E_act_ for step 11 approaching values observed for early-initiation steps 3 and 4 [13].

### Assessing ASA Changes from Disruption of RNAP-Promoter and σ^70^ -Core RNAP interactions in Initiation

Major changes in water-accessible surface area (ΔASA) of RNAP and promoter DNA must occur in the initiation complex before RNAP can escape from the promoter, including the positive ΔASA of disruption of specific RNAP-promoter contacts and release of σ^70^ from core RNAP, as well as the negative ΔASA of duplex formation by the upstream bubble strands. These large-magnitude ASA changes are expected to contribute significantly to the urea and GB kinetic *m*-values of the steps in which they occur. Contributions from each can be estimated from the amount and composition of the ΔASA, using previously-determined solute interaction strengths (α-values; Eq S2) listed in Table S1. No high-resolution structure of an *E. coli* RNAP initiation complex is available for this purpose but the necessary ASA information can be obtained from the published structure of a stable *E. coli* RNAP-λP_R_ promoter open complex (7MKD, thought to be RP_O_ [22]), as described in SI Methods. While the intermediate open complex I_3_ (thought to be that determined as 7MKE [22]) is the species that binds the initiating NTP [14], RNAP - promoter DNA interactions are better-resolved in 7MKD than in 7MKE, allowing ASA calculations to be performed. Initiation complexes at the λP_R_ promoter are very stable [e.g. 71] so it is plausible to model them using the RP_O_ structure. Tables S2-S5 summarize the results of these calculations and the predicted contributions from disruption of RNAP-promoter and σ^70^-core RNAP interactions to urea and GB *m*-values.

### “*Excess*” *m*-Values and E_act_ Values Reveal that RNAP-Promoter Contacts in the Initiation Complex Are Broken In Both Mid- and Late-Initiation Steps and Not Concertedly at the Escape Step

We assume that kinetic *m-*values and E_act_ values of initiation steps 3 and 4 are determined primarily by the ASA and enthalpy changes, respectively, that result from downstream base-pair opening, NTP binding, TL folding and catalysis. These processes also occur in all subsequent steps of initiation (steps 5-11), and we assume that their contributions to ΔASA, *m-*values and E_act_ values are similar for all steps. In addition, steps 5-11 include the processes involved in promoter escape (disruption of RNAP-promoter and σ^70^ - core RNAP interactions, duplex formation by the upstream initiation bubble) which are the main focus of the present study.

We therefore propose that differences in kinetic *m*-values and in E_act_ values between mid- or late-initiation steps (steps 5-11) and earlier steps (steps 3 and 4), which we call *excess* quantities, can be interpreted as the contributions to these kinetic *m*-values and E_act_ values from the processes involved in promoter escape. *Excess* kinetic *m*-values for initiation steps 5-11 are listed in **Table 1** and plotted in **Figure 6**. *Excess* E_act_ values (calculated analogously from previously published results [13]) are also listed in **Table 1** and plotted in **Figure 6**.

**Figure 6.**
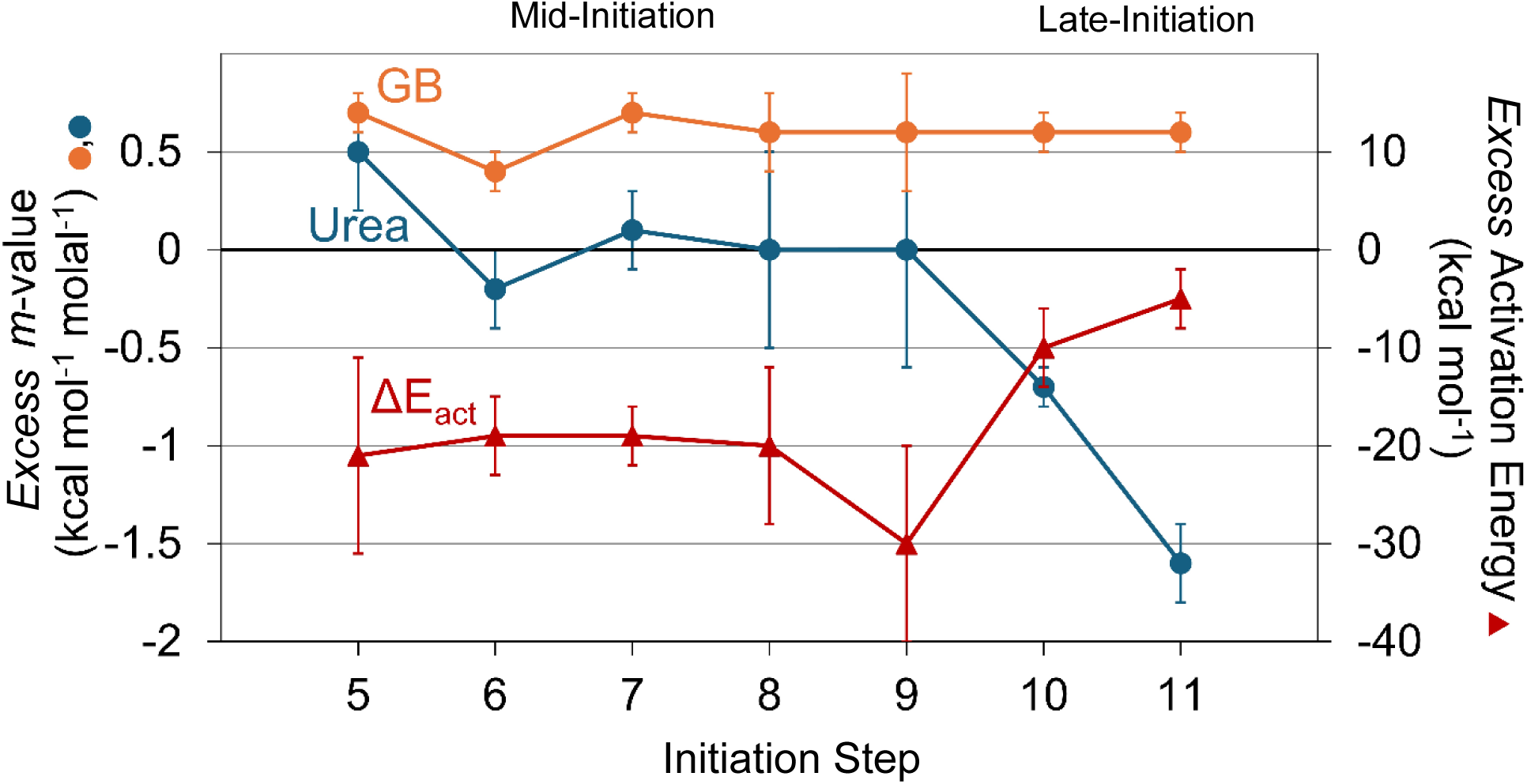
*Excess* Urea and GB Kinetic *m*-values and *Excess* Activation Energies for Mid- and Late-Initiation Steps at λP_R_ Promoter. *Excess* kinetic *m*-values (left axis) and activation energies (right axis), from **Table 1**, are plotted for Mid- and Late-Initiation steps. These *excess* values are defined as differences from those of Early-Initiation steps (steps 3 and 4) that also involve translocation. RNAP-λP_R_ promoter contacts are proposed to break, the upstream duplex is proposed to re-form, and contacts of the σ^70^ specificity subunit with core RNAP are proposed to break in these Mid- and Late-Initiation steps [13, 14, 65].

**Figure 6** shows that moderate- to large-magnitude GB *excess m*-values and *excess* E_act_ values are definitely not confined to the promoter escape step but rather are distributed over many mid- and late-initiation steps. These findings therefore strongly support a step-by-step model rather than a concerted model for disruption of interactions and bubble collapse in the initiation complex. On the other hand, the pattern of urea *excess m*-values (**Figure 6**), which are near-zero for most mid-initiation steps and favorable for late steps, is consistent with a concerted model. However, this pattern can also be explained by a stepwise model if there are compensating contributions to the *excess* urea *m*-values of mid-initiation steps from processes with positive and negative ΔASA. Since GB *excess m*-values and *excess* E_A_ cannot be explained using the concerted model, interpretation of urea *excess m*-values for mid-initiation steps in terms of compensating effects is justified. The quantitative ASA-based predictions of contributions to *excess m*-values from contact disruption and bubble collapse/duplex formation in the following sections support these qualitative conclusions.

### Sums of Urea and GB *Excess m*-Values and E_act_ Values for Mid- and Late-Initiation Steps Agree with Predictions for RNAP-Promoter Contact Disruption, Bubble Collapse, and σ^70^ Release

The logical starting point of a quantitative interpretation of the *excess* quantities in **Figure 6** is to compare their sums over all initiation steps (**Table 1**) with the corresponding sums of predicted contributions from the known events of promoter escape (i.e. disruption of RNAP-promoter and σ^70^ - core contacts and duplex formation by the upstream bubble strands). These predictions, summarized in **Table 2** and described below, agree well with the observed *excess m*-values and E_act_ values for combined steps 5-11.

**Table 2.** Predicted Contributions to Urea and GB m-Values from Disruption of RNAP-DNA and RNAP-RNAP Interfaces and Duplex Formation In Initiation.

| Process | $\Delta ASA$<br>($\text{\AA}^2$) | Urea *m*-value<br>($\text{kcal mol}^{-1} \text{molal}^{-1}$) | GB *m*-value<br>($\text{kcal mol}^{-1} \text{molal}^{-1}$) |
| --- | --- | --- | --- |
| Disrupting $\sigma^{70}$ Contacts with ssDNA (-1 to -11) | $2.9 \times 10^3$ | $-0.8 \pm 0.1$ | $+0.2 \pm 0.3$ |
| Disrupting Core RNAP Contacts with ssDNA | $3.6 \times 10^3$ | $-0.9 \pm 0.1$ | $+1.2 \pm 0.4$ |
| 12-mer Duplex Formation from Half-Stacked Strands | -- | $+1.2 \pm 0.4$ | $+0.3 \pm 0.1$ |
| Sum of Contributions from Disrupting Contacts with ssDNA and Duplex Formation | | $-0.5 \pm 0.4^a$ | $+1.7 \pm 0.5^b$ |
| Sum of *Excess* *m*-values for Steps 5-9 | | $+0.4 \pm 0.8$ | $+3.2 \pm 0.4$ |
| Disrupting $\sigma^{70}$ Contacts with dsDNA (-12 to -38) | $2.3 \times 10^3$ | $-0.6 \pm 0.1$ | $+1.4 \pm 0.2$ |
| Disrupting $\sigma^{70}$ Contacts with Core RNAP | $1.0 \times 10^4$ | $-1.6 \pm 0.3$ | $+1.8 \pm 1.3$ |
| Sum of Contributions from Disrupting $\sigma^{70}$ Contacts with dsDNA and with Core RNAP | | $-2.2 \pm 0.3^c$ | $+3.2 \pm 1.3^d$ |
| Sum of *Excess* *m*-values for Steps 10 and 11 | | $-2.3 \pm 0.2$ | $+1.2 \pm 0.1$ |
| Sum of All Predicted Contributions | | $-2.7 \pm 0.5^e$ | $+4.9 \pm 1.7^f$ |
| Sum of All Observed *Excess* *m*-Values | | $-1.9 \pm 0.8$ | $+4.4 \pm 0.5$ |

<sup>a,c,e</sup> Predicted urea kinetic *m*-value agrees with observed within the combined uncertainties
<sup>b,d</sup> Predicted GB kinetic *m*-value differs from observed by more than the combined uncertainties
<sup>f</sup> Predicted GB kinetic *m*-value agrees with observed within the combined uncertainties

GB *excess* kinetic *m*-values for steps 5-11 (**Figure 6**) are positive and of similar magnitude and sum to 4.4 ± 0.5 kcal mol^-1^ molal^-1^. This experimental result agrees well with that (4.9 ± 1.4 kcal mol^-1^ molal^-1^) predicted from the changes in water-accessible surface area (ΔASA) calculated using the stable λP_R_ RP_O_ structure (**Tables S2-S5**) for the interfaces disrupted and conformational changes that occur before or during escape of RNAP from the promoter. These predictions, summarized in **Table 2**, use solute-atom interaction coefficients obtained previously from model compound studies which are summarized in **Table S1**. The good agreement between predicted and observed quantities provides support for our focus on *excess m*-values and on the interfaces and conformational changes in initiation listed in **Table 2**. Likewise, the sum of predicted urea *m*-value contributions from RNAP-promoter DNA contact disruption, duplex formation and σ^70^ release to the urea *m*-value (−2.7 ± 0.5 kcal mol^-1^ molal^-1^; **Table 2**) agrees with the sum of the experimentally-determined *excess m*-values of these steps (−1.9 ± 0.8; **Table 1**) within the moderately large uncertainties in both values.

*Excess* E_act_ values for steps 5-9 (**Table 1**; **Figure 6**) are all very negative, ranging from −19 to −30 kcal mol^-1^ with uncertainties of ∼30-50%. For step 10 the *excess* E_act_ is −10 ± 7 kcal mol^-1^ while for step 11 it is zero within uncertainty. These sum to −119 ± 16 kcal mol^-1^, which is the enthalpy change expected for formation of 12 stacking interactions in making an internal 11 base pair duplex from half-stacked strands near room temperature [72]. Here also the agreement between observed and predicted values is good. Disruption of contacts between RNAP and promoter strands, especially taking bases out of aromatic pockets on σ^70^, could contribute in the opposite direction, though this contribution may not exceed the uncertainties in the enthalpies used in this comparison. In the following sections, analyses of subsets of *excess m*-values and activation energies for mid- and late steps of initiation are presented.

### ASA Increases from Disruption of σ^70^ Contacts with Upstream Duplex DNA and Core RNAP in Late Initiation Steps Explain the Large-Magnitude, Negative Urea *Excess* Kinetic m-Value of these Steps

The urea *excess* kinetic m-value for late-initiation steps 10 and 11 is −2.3 ± 0.2 kcal mol^-1^ molal^-1^ (**Table 2**). Disruption of contacts between σ^70^ regions 3,4 and the upstream promoter duplex (−12 to −38, including the extended −10 and −35 regions) exposes 2.3 x 10^3^ Å^2^ of surface area (**Table S2**) and is predicted to contribute approximately −0.6 kcal mol^-1^ molal^-1^ to the summed urea *excess* kinetic *m*-value of these two late steps, not nearly large enough in magnitude to explain the observed value. However, inclusion of the effect of disrupting the extensive σ^70^–core interface (1.0 x 10^4^ Å^2^), with a predicted *m*-value contribution of −1.6 ± 0.3 kcal mol^-1^ molal^-1^ (Tables 2, S6) brings predicted and observed values into agreement. While some interactions of σ^70^ region 3.2 with core RNAP are found to be disrupted earlier in mid-initiation steps [e.g. 23], we estimate that these involve less than 10% of the total ASA exposed when σ^70^ is released from the initiation complex and therefore should not greatly affect the ASA-based analysis. Insufficient structural information is available to include interactions of the αCTD with the λP_R_ UP element in this analysis, but this may not matter because the λP_R_ UP element deviates significantly from consensus [73] and nonspecific interactions of the αCTD with upstream DNA may persist after those with the UP element are disrupted.

### The Large Negative Urea *Excess m*-Value and Modest *Excess* Activation Energy of Late Steps of Initiation Indicate that the Upstream Bubble Does Not Convert to Duplex in these Steps

In a concerted model of contact disruption and promoter escape, collapse of the upstream initiation bubble and duplex formation would occur together with σ^70^ release late in initiation. But if the contribution to the urea *excess m*-value from duplex formation from separated, half-stacked strands (+1.2 ± 0.4 kcal mol^-1^ molal^-1^) is included with those predicted above for disruption of σ^70^ contacts with upstream duplex DNA and core RNAP in these late steps, the resulting predicted urea *excess m*-value (−1.0 ± 0.5 kcal mol^-1^ molal^-1^) differs from the observed value (−2.3 ± 0.2 kcal mol^-1^ molal^-1^) by more than the combined uncertainties. By contrast, including the contribution from duplex formation in the prediction of the urea *m*-value for mid-initiation steps improves the agreement with the observed value, as described below. These findings are completely consistent with the conclusion drawn from the *excess* enthalpy profile in **Figure 6** (see also [13]). The finding of large-magnitude negative *excess* enthalpies only in steps 5–9 is completely consistent with a step-by-step process of contact disruption and duplex formation in these mid-initiation steps. The alternative model of concerted duplex formation in RNAP escape would mean this entire negative enthalpy (approximately −120 kcal) would contribute only to the E_act_ value(s) for the last step or two of initiation, which from **Figure 6** is clearly not the case.

### The Small-Magnitude *Excess* Urea Kinetic m-Value and Negative *Excess* Activation Energies of Steps 5-9 Are Consistent with Models in which Bubble Collapse and Duplex Formation Occur As Contacts Between RNAP and Bubble DNA Strands are Broken in these Steps

Both σ^70^ and core RNAP have significant interactions with the strands of the upstream initiation bubble (−1 to −11). Disrupting the interface between σ^70^ (regions 1.2 and 2) and the discriminator and −10 regions of the nt strand exposes 3.6 x 10^3^ Å^2^ of surface area (**Table S3**). If this occurs in steps 5-9, as proposed [13, 14] it will contribute −0.8 ± 0.1 kcal mol^-1^ molal^-1^ to the summed urea *excess* m-value of these steps (**Table S3**). Likewise, disrupting the interface between core RNAP and the t strand of the initiation bubble exposes 2.9 x 10^3^ Å^2^ of surface area (**Table S4**). If this occurs in steps 5-9, it will contribute approximately −0.9 kcal mol^-1^ molal^-1^ to the summed urea *excess* m-value of these steps (**Table S4**). The sum of these contributions (−1.7 ± 0.1 kcal mol^-1^ molal^-1^) is significantly larger in magnitude than the sum of urea *excess* kinetic m-values for these steps (+0.4 ± 0.9). Much if not all of this difference can be explained if duplex formation by the bubble strands (predicted urea *m*-value of 1.2 ± 0.4 for initially half-stacked strands [72, 74]) accompanies the disruption of their interactions with σ^70^ and core RNAP in steps 5-9. Including duplex formation, the urea *excess* kinetic *m*-value is predicted to be −0.5 ± 0.4 kcal mol^-1^ molal^-1^, which is small in magnitude like the observed value (+0.4 ± 0.9 kcal mol^-1^ molal^-1^; **Table 2**) and not significant different from it, given the uncertainties in both quantities. Stepwise duplex formation from half-stacked strands, with an enthalpy change of approximately −10 kcal mol^-1^ per base pair formed [72], was proposed previously to explain the E_act_ values of steps 5-9 [13]. No other conformational change that occurs or might occur in these steps of initiation (e.g. scrunching of the strands, movements of σ^70^ region 3 on interacting with the hybrid) appears capable of explaining these *excess* E_act_ values and urea kinetic *m*-values.

### Moderately Large Unfavorable *Excess* GB Kinetic m-Values of All Steps May Indicate a Step-by Step Loosening of RNAP-Promoter Interactions, Particularly with DNA Phosphates, In Mid-Initiation Steps

Predictions for contributions from disruption of the different RNA-DNA and RNAP-RNAP interfaces and from duplex formation to GB *excess* kinetic m-values of steps 5-9 and 10-11 are given in **Table 2**. These predictions, like the observed GB *excess* kinetic *m*-values of these groups of steps, are large and positive, but predictions for the events of steps 5-9 are approximately 1.5 kcal mol^-1^ molal^-1^ less positive than the sum of observed *excess* GB kinetic *m*-values, while predictions for events of steps 10-11 are approximately 2 kcal mol^-1^ molal^-1^ more positive than observed. Since sums of predicted and observed *excess* GB kinetic *m*-values for all steps (5-11) agree within uncertainty (**Table 2**), we interpret the offsetting predicted-observed differences for mid- and late initiation steps as an indication that some contacts between σ^70^ and core RNAP and/or upstream duplex DNA may be disrupted in mid-initiation steps, in a way that greatly affects GB *m*-value predictions but not urea *m*-value predictions. Exposure of carboxylate and phosphate ASA are likely candidates; these atom-types contribute disproportionately to the predicted GB *m*-values compared to urea (**Table S5**). Approximately 12 carboxylates (estimated from 959 Å^2^ at ∼80 Å^2^ per carboxylate) and 7 phosphates (estimated from 523 Å^2^ at ∼75 Å^2^ per phosphate) are exposed/rehydrated in dissociation of σ^70^ from core RNAP and from duplex promoter DNA, respectively, contributing 1.62 ± 0.10 kcal mol^-1^ molal^-1^ and 1.49 ± 0.12 kcal mol^-1^ molal^-1^, respectively, to the predicted GB *m*-values of these dissociations. If half of this rehydration occurs in steps 5-9, perhaps from weakening of the interfaces of σ^70^ regions 1.2, 2 and/or 3 with core RNAP and of the interface of σ^70^ region 3 with the extended −10 region of duplex promoter DNA in the process of disrupting contacts of RNAP with the strands of the discriminator and −10 regions, this could account for the differences between predicted and observed GB *excess* kinetic m-values. The predicted effect of this rehydration of phosphates and carboxylates on the urea *m*-value is only 12% as large in magnitude (Table S1) and would not be detectable given the uncertainties.

Precedent for this proposal is provided by the dissociation of lac repressor protein from lac operator DNA [70]. Mechanistically, dissociation occurs in several steps, the first of which is disruption of the protein-protein interaction of the operator-DBD-hinge helix assembly with the core of lac repressor (DBD is the acronym for the DNA binding domain of lac repressor). Subsequently the hinge helices unfold, and the operator DNA dissociates from the two repressor DBD. Though all the phosphates are of course part of these latter interfaces, all of the GB effect is on the first step. Clearly the disruption of the protein-protein interface loosens the protein-DNA interface to allow rehydration of DNA phosphates prior to DNA dissociation [70]. These hydrated phosphates would still interact coulombically with the DBD.

Another factor that, in principle, could affect the GB *excess* kinetic *m*-value for steps 10-11 is the interaction of two polyanionic (each 15-20 carboxylate) regions of free σ^70^ with its DNA-binding domains [75, 76]. However, although these interactions are strong enough to result in autoinhibition of binding of free σ^70^ to promoter DNA, they are not specific enough to be detected in σ^70^ structures. Therefore, these interactions are probably coulombic interactions between hydrated charged groups and, in the absence of dehydration, would not affect solute *m*-values greatly.

### Comparison of Mechanisms of Contact-Disruption in Initiation and Promoter-Escape by *E. coli* Eσ^70^ RNAP and T7 Phage RNAP

T7 phage RNAP, like *E. coli* Eσ^70^ RNAP, interacts with three regions of promoter DNA upstream of the start site to bind specifically to the promoter and open the start site region. Structural, biochemical and/or kinetic-mechanistic evidence indicate that these interactions are disrupted sequentially from downstream to upstream in three groups of steps as the RNA-DNA hybrid translocate in initiation.

In the complex of T7 RNAP with a 17 bp promoter fragment (−1 to −17) the upstream initiation bubble spans the region −1 to −4 [53]. This region is open because of binding interactions of RNAP side chains with the bubble template strand bases, as well as with the downstream face of the −5 bp. Synthesis of a 3 bp RNA-DNA hybrid, involving one translocation, disrupts the interaction with −1, scrunching this residue, and repositions t-strand residues −2 and −3. Reference is made to another structure obtained in the presence of the next NTP (and therefore translocated an additional step) in which no contacts with t-strand bubble residues −1 to −4 are detected, indicating that their contacts with RNAP are disrupted [53]. Interactions with −5 and upstream bp are preserved until the hybrid extends to 7 or 8 bp [51]. Consistent with this, we [14] concluded from analysis of kinetic data for steps of NTP incorporation by a T7 RNAP initiation complex [35] that translocation free energies for steps synthesizing 3-mer to 5-mer RNA are all relatively unfavorable and that the step synthesizing a 4-mer RNA is the most unfavorable of these steps. Residues −1 to −4 of the bubble nt strand are not resolved in these structures and hence presumably interact less strongly and/or are disordered.

In the *E. coli* Eσ^70^ RNAP-λP_R_ promoter open complex (RP_O_) both strands of the discriminator region (−1 to −6), the downstream portion of the bubble, interact with RNAP to stabilize the open state [22]. From activation energies and translocation free energies for individual steps of initiation by *E. coli* Eσ^70^ RNAP at the λP_R_ promoter, we proposed that contacts with discriminator strand residues −1 to −6 are disrupted in extension of the RNA-DNA hybrid from 3 bp to 6 bp [13, 14]. In the current research, disruption of contacts with discriminator and −10 regions of the upstream bubble are considered together to interpret composite kinetic *m*-values for extension of the RNA-DNA hybrid from 4 bp to 9 bp.

For both T7 and *E. coli* RNAP, interactions with the mid-region (−7 to −11) of the promoter appear to be the primary contributors to OC stability and specificity. For T7 RNAP, these are interactions of the specificity loop with the promoter duplex [36, 37, 39, 51, 53, 54], while for *E. coli* σ^70^ RNAP these are interactions of σ^70^ region 2 and the β and β’ subunits of core RNAP with the individual strands of the open −10 region [22, 24]. For T7 RNAP, multiple lines of evidence indicate that these interactions are broken in extending the RNA-DNA hybrid from 5-mer to 8-mer [37, 39]. In agreement with this, steps that extend the hybrid from 6-mer to 9-mer exhibit a second group of relatively unfavorable translocation free energies, indicating that strong contacts are disrupted in these steps [14]. For *E. coli* RNAP, from the patterns of unfavorable translocation free energies [14] and activation energies [13], we concluded that contacts with the strands of the −10 region are disrupted as the hybrid extends from 6-mer to 9-mer.

Farther upstream, the AT-rich duplex region from −13 to −17 of the T7 consensus promoter is recognized by minor groove interactions with R96, K98 and other residues of a surface loop (residues 93 – 101 [54]). Contacts between the AT-rich promoter region from −13 to −17 with the AT recognition loop of T7 RNAP are disrupted as the hybrid extends from 8-mer to 12- or 13-mer [37, 39] Large unfavorable translocation free energies are calculated for three of these steps (synthesis of 10-, 12-, 13-mer RNA), interpreted as the costs of disrupting these interactions and of accompanying conformational changes [14].

Upstream of the open region of the λP_R_ promoter, the σ^70^ subunit of E coli RNAP makes specific interactions with the extended −10 duplex (−13 to −17) and the −35 region (approximately −33 to −38), while the CTD of the two α-subunits interact with the UP element of the promoter (−40 to −60) and weaker nonspecific interactions occur to approximately −80 [9, 22]. These upstream duplex contacts are inferred to break as the hybrid extends from 9-mer to 11-mer, modestly increasing the translocation cost for 10-mer synthesis and greatly increasing it for 11-mer [14]. For both RNAP, escape from the promoter occurs when these upstream contacts are disrupted, in conjunction with other conformational changes in the transition from initiation to elongation [36, 52, 65].

### Timing of Upstream Bubble Collapse/Duplex Formation

For *E. coli* RNAP at the λP_R_ promoter, urea *excess* kinetic *m*-values and *excess* activation energies obtained from nucleotide-incorporation rate constants (cf. **Figure 6**) are consistent with a step-by-step model of collapse of the upstream bubble (beginning at −1 and ending at −11) and duplex formation, and inconsistent with a concerted model of bubble collapse at the escape point. These activation energies indicate that bubble collapse/duplex formation occurs in five or six steps, each forming ∼2 bp, once contacts of those bases with RNAP are disrupted [13]. The discriminator and −10 strands may convert to duplex in steps 5-7 and 8-9, respectively.

Information about changes in the *E. coli* RNAP initiation bubble as the RNA-DNA hybrid extends is available for one other promoter (T5 N25). Real-time single-molecule torsional force studies with this promoter during RNAP binding, OC formation, productive initiation and escape revealed that prior to promoter escape the amount of unwound (bubble) DNA in the initiation complex increased to approximately twice that of the binary OC [20]. This result indicates that bubble collapse and duplex formation at T5 N25 occur late in initiation.

One possible explanation for the different timing of bubble collapse in initiation complexes at λP_R_ and T5 N25 promoters involves the likely difference in strength of interactions of downstream mobile elements (DME, including the β lobe and β’ jaw and SI3 domains [77, 78]) with the downstream duplex (DD, extending from +6 to +20). The strength of these DME-DD interactions is determined allosterically by the strength of the “clasp” structure, which reinforces interactions between the non-template strand of the upstream discriminator (−6 and −5 for λP_R_ and T5N25) and the RNAP cleft (σ^70^ regions 1.1 and 1.2 and the β-lobe). In the λP_R_ OC, these interactions promote a closed clasp conformation, reinforcing contacts between the σ^70^ 1.1-linker and the β gate loop and causing the ejection of σ^70^ region 1.1 from the downstream channel.

These conformational changes align the downstream duplex DNA tightly within the channel, promoting strong downstream contacts [4, 22, 79, 80]. G residues at the first and second positions from the upstream end of the discriminator nt-strand result in a longer-lived OC than for other nucleotides investigated [22, 81, 82]. Structurally, comparison of λP_R_ with λP_R_ −5C reveals that λP_R_ has a tighter clasp and stronger DME-DD interaction [22]. Consistent with this, the lifetime of the WT λP_R_ RP_O_ complex (with GG at −5, −6 positions) exceeds that of the mutant λP_R_ −5C open complex by 4- to 5-fold at the same temperature and salt concentration [22]. This lifetime difference is accompanied by a large structural difference. The OC at mutant λP_R_ −5C promoter lacks many of the strong σ^70^ region 1.2-discriminator and DME-downstream duplex interactions of λP_R_ RP_O_ [22]. T5 N25 (with AG at −5, −6) differs from WT λP_R_ at the −6 position, which from the structure is also involved in many specific interactions with σ^70^ region 1.2 [22], and exhibits an OC lifetime [83] which is only 30% that of λP_R_ RP_O_ at the same conditions. Though structural information about the T5 N25 OC is not available, it is likely that it is similar to λP_R_ −5C and lacks the strong σ^70^ region 1.2-discriminator and DME-downstream duplex interactions of λP_R_.

This difference in DME-DD interactions between λP_R_ and T5 N25 promoters probably persists at least until synthesis of a 5 or 6 RNA-DNA hybrid, when interactions of σ region 1.2 with upstream discriminator bases are thought to be disrupted as discussed above. Strong DME-DD interactions could limit the ability of DD to rotate when a bp is opened at the active site in translocation, introducing the possibility that the twist generated by opening each downstream bp is fed upstream in translocation and utilized to form an upstream bp once the contacts of these bases with RNAP that favored bubble formation initially are broken. Indirect support for the hypothesis that strong DME-DD interactions limit the ability of DD to rotate is provided by the observation that these strong DME-DD interactions aren’t present in I_2_, the unstable intermediate OC, and form only after opening of the initiation bubble is complete [6, 7]. T5 N25 and λP_R_ sequences of course differ at other positions but the difference in the upstream discriminator (AG vs GG) appears most likely to explain the difference in OC lifetime and the inferred difference in strength of DME-DD interactions which, in turn, might explain the differences between these two promoters in the timing of bubble collapse in initiation complexes.

Bubble collapse and duplex formation occur late in initiation by T7 phage RNAP [39] and by eukaryotic Pol II RNAP in complexes with multiple upstream and downstream transcription factors [55, 56]. T7 RNAP lacks a DME apparatus analogous to *E. coli* RNAP to constrain rotation of the downstream duplex, eliminating any mechanism like that proposed here to allow duplex formation by the upstream bubble strands to occur prior to the escape point [39]. Pol II has an extensive downstream apparatus including TFIIH, a helicase that drives translocation and rotation of the downstream duplex DNA while the upstream duplex is immobilized by binding interactions [55, 56]. TFIIH thereby opens the initiation bubble and also drives each step of translocation and rotation involved in opening a downstream bp in initiation. These actions of TFIIH may prevent bubble collapse from occurring before the escape point for pol II.

### Conclusions

We report the large effects of urea and glycine betaine on individual steps of nucleotide incorporation into the growing RNA-DNA hybrid in transcription initiation and escape of RNA polymerase from promoter DNA. These solute effects are quantified as kinetic *m*-values, as in previous studies of mechanisms of protein folding [69] and formation of a repressor-operator complex [70]. We conclude from analysis of these urea and GB kinetic *m*-values that RNAP-λP_R_ promoter contacts are not broken concertedly at the escape point but rather break in multiple mid- and late steps of initiation, as previously deduced from activation energies and magnitudes of these rate constants [13, 14]. Our solute studies support the previous finding [17, 65] that σ^70^ is released in promoter escape and support our previous conclusion [13] that at the λP_R_ promoter the upstream duplex does not re-form concertedly at the escape point but rather forms in multiple mid-steps of initiation once upstream RNAP-promoter contacts are broken in these steps.

## Materials and Methods

Methods of sample preparation, fast kinetics studies of transcription initiation, gel electrophoretic separations of RNA products, phosphoimager analysis, and interpretation using ASA analysis of high resolution PDB structures were largely as described previously. Changes to these published methods are described in SI Methods.

## Acknowledgements.

This research was supported in part by NIH GM R35 118100 and also by the UW Madison Graduate School and Department of Biochemistry.

