## Supplemental Methods and Data for "Evidence for Stepwise Disruption of *E. coli* RNA Polymerase - λP_R_ Promoter Contacts and Bubble Collapse in Transcription Initiation"

**Departments of Biochemistry[1] and Chemistry[3] and Biophysics Program[2]**

**University of Wisconsin Madison**

**Madison WI 53706**

### Materials and Methods

#### Reagents, Buffers, Gels

Reagents used in buffers and stock solutions were purchased in the highest grade available and used as received. All solutions were prepared using 18 MΩ deionized water and filtered. Stock solutions of urea (10 M, made with Ultrapure (> 99%) urea from DOT Scientific) and glycine betaine (GB) (5 M, made with > 99% anhydrous GB from Sigma-Aldrich) were used in transcription experiments. NTPs for transcription assays and dNTPs for PCR reactions (all > 99%, from Thermo Fisher Scientific, Waltham, MA) were used as received. Enzymes for PCR reactions were from New England Biolabs (NEB, Ipswich, MA) and used according to the manufacturer’s protocols.

RNAP storage buffer (SB) is 50% v/v glycerol, 0.01 M Tris, 0.1 M NaCl, 0.1 mM EDTA, and 0.1 mM DTT. Transcription buffer (TB) is 40 mM Tris (pH 8.0), 5 mM $MgCl_2$, 60 mM KCl, 1 mM DTT, and 0.05 mg/mL BSA. "High UTP" (also low GTP) initiation solution (IS) used in most experiments is 400 μM ATP, 400 μM UTP, 20 μM GTP, 70 nM α-$^{32}$P GTP and 100 μg/mL heparin in TB. Other experiments were performed with a "low UTP" (also high GTP) IS that is 400 μM ATP, 400 μM GTP, 20 μM UTP, 70 nM α-$^{32}$P UTP and 100 μg/mL heparin in TB. Urea or GB is added to IS at twice the desired final concentration (≤ 2 M urea; ≤ 3 M GB). Quench solution (QS) is 8 M urea and 15 mM EDTA in TB. Loading dye is 0.05% w/v xylene cyanol and 0.05% bromophenol blue in QS.

All transcription gels are 20% acrylamide-(bis)acrylamide (19:1), 7.5 M urea and are made using the UreaGel system (National Diagnostics).

**Preparation of Promoter DNA and RNA Polymerase**

$\lambda P_R$ promoter DNA, embedded in a 124 bp DNA fragment (-82 to +42) was prepared by PCR as described previously [1]. Briefly, oligonucleotides with an overlapping region are annealed and filled with DNA polymerase, and these filled-in primers are amplified via PCR and purified. As in previous studies [1-4], the initial transcribed region (ITR) of $\lambda P_R$ is modified so that all encoded C positions in the template sequence prior to position +17 are changed to G.

*E. coli* RNAP core enzyme and $\sigma^{70}$ subunit were overexpressed and purified as described previously [1]. To form RNAP holoenzyme, the core enzyme is incubated at a 2:1 mole ratio with $\sigma^{70}$ in SB for 1 hour at 37 °C, then stored at -20 °C. RNAP concentrations reported here are for RNAP holoenzyme that is "active" in open complex (OC) formation at the $\lambda P_R$ promoter at 37 °C, determined by filter binding with a heparin challenge [2]. In the RNAP preparation used here, approximately 50% of RNAP holoenzyme molecules were active in OC formation.

**Transcription Assays, Gel Separations of $^{32}$P-labeled RNAs, and Phosphorimager Quantification**

Fast kinetic studies of transcription initiation were performed using a Kintek Rapid Quench Flow (RQF) instrument as described previously [2-4] with minor modification as below. OC were formed by incubating RNAP holoenzyme and $\lambda P_R$ promoter DNA (final concentrations of 200 nM and 100 nM, respectively) in TB for 1 hr at 19 °C. 20 μL of a solution of preformed OC (final concentration 50 nM) was combined with 20 μL IS (final NTP concentrations half as large as in IS) using the RQF mixer, allowed to react, and then rapidly quenched at 14 different times (from 0.1 s to 120 s) with a 10-fold excess of QS. At least two independent series of experiments were performed at each NTP condition and solute concentration investigated. Additional experiments in the absence of urea and GB were performed at both NTP conditions investigated as controls and for comparison with previously-published results [2].

An aliquot (8.8 µL) of each quenched timepoint was combined with loading dye (1.2 µL) and 5 µL of the mixture was loaded onto a 20% denaturing PAGE gel and separated at ~2 kV for 3-5 hours. PAGE gels were imaged with an Amersham Typhoon and analyzed with ImageQuant software as described previously [2].

Kinetic analysis of phosphoimager results was performed as previously described [2] to obtain RNA population fractions as a function of time for all lengths, except that all experiments at a given NTP and solute concentration were fitted together to obtain rate constants, rather than fitting them individually and averaging the results. To fit results of multiple experiments together, amounts of RNA transients as functions of time in each experiment were normalized by the final amount of full length (11+) RNA in that experiment.

**Calculations of Changes in Water Accessible Surface Area ($\Delta$ASA) From Disruption of $\sigma^{70}$-DNA, Core-DNA and $\sigma^{70}$-Core Contacts in Transcription Initiation from Structural Information**

ASA calculations were performed using the program SurfRacer [5, 6] with a water-probe radius of 1.4 Å on the high-resolution cryoEM structure of the stable OC ($RP_O$) formed by E. coli RNAP holoenzyme with a 90 bp $\lambda P_R$ promoter construct (-60 to +30; 7MKD [7]). The DNA is resolved from approximately -55 to +23, including most of the UP element. Only the proximal $\alpha$CTD is observed as bound in 7MKD. To determine the amount and composition of the $\Delta$ASA from disrupting $\sigma^{70}$-DNA contacts, the core subunits were deleted from the $RP_O$ structure and the ASA of the resulting $\sigma^{70}$-DNA complex was compared with the ASA of $\sigma^{70}$ (with DNA deleted) and DNA (with $\sigma^{70}$ deleted). This $\Delta$ASA information was divided into contributions from disrupting contacts of $\sigma^{70}$ with the bubble strands and contacts with the upstream duplex regions (extended -10, -35) by assuming that regions 1.2 and 2 of $\sigma^{70}$ (ending at residue 455) contact the bubble strands (-11 to -1) and that regions 3 and 4 of $\sigma^{70}$ (starting at residue 456) contact the duplex DNA upstream of -11.

To determine the amount and composition of the $\Delta$ASA from disrupting core RNAP contacts with the bubble strands, the $\sigma^{70}$ subunit was deleted from the $RP_O$ structure and the ASA of the resulting core RNAP-DNA complex was compared with the ASA of core (with the bubble strands of DNA deleted) and the bubble DNA (with core deleted). The only specific contacts of core RNAP with the duplex region of the promoter above -11 are interactions of the $\alpha$CTD subunits with the UP element. ASA analysis revealed that the contributions to overall urea and GB m-values for initiation (Table 1) from disruption of the subset of UP element interactions with the single resolved α-CTD in 7MKD were too small in magnitude to be significant and were therefore neglected.

To determine the amount and composition of the ΔASA from disrupting $\sigma^{70}$-core contacts the promoter DNA was deleted from the $RP_O$ structure, and the ASA of the resulting $\sigma^{70}$-core complex was compared with the ASA of $\sigma^{70}$ (with core deleted) and core (with $\sigma^{70}$ deleted).

**Calculations of Changes in Water Accessible Surface Area (ΔASA) for Binding the Initiating NTP and Trigger Loop (TL) Folding in RNA Dinucleotide (pppApU) Synthesis from Structural Information**

Because the RQF experiments reported here provide no information about binding of the initiating ATP and UTP, TL folding and the kinetics of dinucleotide (pppApU) synthesis as a function of solute concentration, *m*-value predictions for these steps from structural information were used in the global fitting of RQF data to obtain *m*-values for subsequent steps as described elsewhere in SI. No high-resolution structural information is yet available for these first steps in transcription initiation by *E. coli* RNAP. Therefore, the high-resolution crystal structure of a *T. thermophilus* initiation complex with ATP and a nonhydrolyzable CTP analog (CMPCPP) bound in the active site (4Q4Z; [8]) was used for ASA calculations with SurfRacer [5, 6]. CMPCPP has phosphorus-carbon bonds (P-C-P) between the alpha and beta phosphates instead of the phosphorus-oxygen (P-O-P) bonding in CTP. Because the ASA of this position in the triphosphate group is negligibly small, this change does not affect the ASA analysis. ATP and UTP are the initiating nucleotides at the $\lambda P_R$ promoter. UTP differs from CTP in the substituent on the C1 position of the base, namely a carbonyl ($sp^2$) oxygen for UTP vs an amino ($sp^3$) nitrogen for CTP. Predictions from an ASA analysis using the α-values of Table S1 reveals that, even if the substituent is completely buried in binding to RNAP, substitution of UTP for CTP would affect *m*-value predictions by only ~8 cal $mol^{-1}$ $molal^{-1}$ for urea and ~100 cal $mol^{-1}$ $molal^{-1}$ for GB. These predicted maximum differences between CTP and UTP are smaller than the uncertainties in the predicted urea and GB *m*-values for CMPCPP binding (340 ± 30 cal $mol^{-1}$

molal$^{-1}$ for urea, - 880 $\pm$ 150 cal mol$^{-1}$ molal$^{-1}$ for GB; see Table S7) and therefore not significant.

Seven residues (1246-1251) of the important, conformationally-variable trigger loop (TL) - trigger $\alpha$-helix (TH) region are missing from the 4Q4Z structure with bound ATP and CMPCPP, presumably because they are unfolded. In the absence of NTP, the entire TH/TL region is absent from crystal structures of OC formed with *T. Th.* RNAP (4G7H, residues 1238-1251; [9]) and *E. coli* RNAP (7MKD, residues 1127-1134; [7]), indicating that this region is unfolded before binding the initiating NTP. Both nucleotides were deleted to model the initial state of the RNAP-promoter complex before nucleotide binding. To model the initiation complex with only ATP bound, CMPCPP was deleted from 4Q4Z. TL residues 1219-1245 and 1253-1266 convert to TH on binding the active-site NTP [10-12]. The *T. thermophilus* amino acid sequences of these unfolded regions of the TL were built into the structures used for ASA analysis as flexible chains using PyMOL [13] and refined manually with the Regularize Zones tool in WinCoot [14].

These structures and those of the unbound nucleotides (also from 4Q4Z) were analyzed using SurfRacer [5, 6] to obtain amounts and compositions of the $\Delta$ASA for a) binding the first initiating NTP to an RNAP-promoter OC with an unfolded TL, b) binding the second initiating NTP to this complex, accompanied by partial folding of the TH, and c) folding the remainder of the TH as part of catalysis of phosphodiester bond formation. Steps b) and c) do not account for possible movements or conformational changes in downstream mobile elements like SI3 and the downstream mobile jaw that may be coupled to TH formation. This analysis may therefore account for only part of the ASA changes that occur in the overall process of incorporating the 2$^{nd}$ nucleotide into the growing RNA chain.

Results of these analyses are listed in SI Tables S6-S9. Table S6 shows that binding of the initiating ATP buries 402 Å$^2$ of ATP ASA and 217 Å$^2$ of RNAP ASA. Using the urea and GB α-values listed in Table S1, urea and GB *m*-values for ATP binding are readily predicted. From

Table S6, urea disfavors ATP binding moderately (0.21 $\pm$ 0.02 kcal $mol^{-1}$ $molal^{-1}$) while GB favors ATP binding to a similar extent (- 0.21 $\pm$ 0.06 kcal $mol^{-1}$ $molal^{-1}$). Phosphate oxygens are predicted to make the largest contributions to urea and GB *m*-values for ATP binding because they contribute more than 20% of the ΔASA (Table S6) and the urea and especially GB α-values for phosphate oxygen (Table S1) are relatively large in magnitude.

Table S7 shows that binding of the second initiating nucleotide buries twice as much ASA as for binding the first NTP: 494 $Å^2$ of CMPCPP and ATP ASA and 851 $Å^2$ of RNAP ASA. The much larger-magnitude ΔASA contribution from RNAP predicted for CMPCPP binding than for ATP binding arises primarily from TL folding to TH and TH-nucleotide interactions. Using the urea and GB α-values listed in Table S1, urea and GB *m*-values for CMPCPP binding are readily predicted (Table S7). Urea is predicted to disfavor CMPCPP binding moderately (0.34 $\pm$ 0.03 kcal $mol^{-1}$ $molal^{-1}$), while GB greatly favors CMPCPP binding (- 0.88 $\pm$ 0.15 kcal $mol^{-1}$ $molal^{-1}$). This much larger-magnitude predicted GB m-value for binding the second initiating nucleotide as compared to the first results primarily from larger-magnitude ΔASA for phosphate, amide and carboxylate oxygens, all of which interact very unfavorably with GB when accessible.

After rapidly reversible binding of the two initiating nucleotides, the rate-limiting step for dinucleotide synthesis is thought to be folding of the remainder of TL (residues 1246-1252) to enclose the active site step, prior to the faster chemical reaction of NMP addition [12, 15]. Table S8 summarizes the changes in RNAP and DNA ASA in this step, both of which are modest because only seven RNAP residues fold, changing their exposure and that of DNA phosphates in the active site.

The combination of these ASA and *m*-value analyses (Table S9) predicts that the overall step of binding the second nucleotide and synthesis of the initial dinucleotide is moderately disfavored by urea (0.46 ± 0.04 kcal/mol/molal) and significantly favored by GB (-0.97 ± 0.16

kcal/mol/molal). The large-magnitude GB m-value predicted for this first step of RNA synthesis arises primarily from burial of nucleotide phosphate oxygens, with a contribution (-0.78 ± 0.05 kcal/mol/molal) that is almost 80% as large as the predicted GB m-value.

**Data Analysis**

Two or three complete sets of kinetic determinations were obtained for each solute concentration and NTP condition investigated (high and low UTP for urea; high UTP for GB), as exemplified by **Figures 1-2** and **S1-2**. Two types of analysis of these kinetic data were performed. For the kinetic *m*-value plots in **Figure 3**, initiation kinetic data for productive complexes at one solute concentration were fit to the 11-step mechanism of initiation as previously described [2] to obtain rate constants for each initiation step at that solute concentration. To obtain more accurate *m*-values for steps 3-11, global fitting of all kinetic data at all concentrations of each solute was performed with Kintek Explorer by introducing **Eq. 6** for the relationship between rate constants for each step of initiation at different solute concentrations and the solute-concentration-independent *m*-values of these step.

Rate constants for binding and dissociating the two initiating NTP and for catalysis of pppApU synthesis at the chosen solute concentration are needed for both analyses of the kinetic data. These rate constants were calculated from previously-published values in the absence of solute [2, 3] using the predicted solutes *m*-values in SI Tables S6-8. These predictions are thermodynamic *m*-values quantifying the dependences of NTP equilibrium binding constants on solute concentration. Because individual forward and reverse rate constants for these initial NTP binding steps are used in the kinetic fitting, three separations of the thermodynamic *m*-value were tested, in which either 0%, 50% or 100% of the solute effect is in the NTP dissociation rate constant. Putting most if not all of the solute effect in the dissociation rate constant is justified if the association kinetics are diffusion-limited, as may be the case [16, 17]. For TL folding, where only the forward kinetics are assumed to be relevant provided this is the rate determining step [12, 15], the same range (0-100% of the predicted thermodynamic *m*-value) was tested. Similar fits, not significantly different given the uncertainties, were obtained for these various situations.

Predicted urea and GB kinetic *m*-values for TL folding in pppApU synthesis (Table S8) are 0.10 $\pm$ 0.01 and - 0.13 $\pm$ 0.05 kcal $mol^{-1}$ $molal^{-1}$ respectively. Reducing these kinetic *m*-values had no significant effect on fitted kinetic *m*-values for subsequent steps. Increasing the urea kinetic *m*-value for pppApU synthesis more than fourfold (to 0.46 kcal $mol^{-1}$ $molal^{-1}$) also had no significant effect on the fit, while increasing it ten-fold (to 1.03 kcal $mol^{-1}$ $molal^{-1}$) reduced the 3-mer kinetic *m*-value by a marginally-significant amount without affecting kinetic *m*-values for other steps. Likewise, increasing the GB kinetic *m*-value for pppApU synthesis from -0.13 $\pm$ 0.05 kcal $mol^{-1}$ $molal^{-1}$ to 0.11 or 0.49 kcal $mol^{-1}$ $molal^{-1}$ had small, marginally-significant effects only on fitted kinetic *m*-values for 3-mer and 4-mer, without affecting kinetic *m*-values for other steps. This analysis shows that the globally-fitted kinetic *m*-values in Table 1 for steps 3-11 are robust. Only a massive conformational change in addition to TL folding accompanying pppApU synthesis, for which there is no precedent as discussed above, could affect the *m*-values estimated for these steps sufficiently to modify those obtained from the fitting for subsequent steps.

**Background on How Urea and Glycine Betaine Affect Biopolymer Processes and on the Analysis of these Effects**

Urea interacts favorably with most types of unified (i.e, with bonded H atoms) O, N and C atoms of proteins, nucleic acids, and model compounds – especially amide and carbonyl oxygens, aromatic carbons and nitrogens [18-21]. The thermodynamics of these short-range urea-atom interactions and their consequences for biopolymer processes have been quantified, and in some cases dissected, into atom-atom interactions [18-24]. The thermodynamic contribution of interactions of urea with a given type of O, N or C atom increases in proportion to the water-accessible surface area (ASA) of that atom, with a proportionality constant ($\alpha$-value) that quantifies the strength of that interaction per unit ASA. These contributions from interactions of urea with different atom-types are additive, as indicted below. Urea destabilizes folded proteins, $\alpha$-helices and nucleic acid helices because of its favorable interactions with the O, N and C ASA exposed in unfolding (denaturing) these structures (i.e. the $\Delta$ASA) [18-21].

Interactions of GB with O, N and C unified atoms of model compounds and biopolymers are described and quantified as for urea above. Unlike the situation for urea, GB-atom interactions range from quite favorable to strongly unfavorable. GB interacts favorably with cationic and amide nitrogens as well as aromatic ($sp^2$) carbons but interacts slightly unfavorably with aliphatic ($sp^3$) carbon and very unfavorably with DNA phosphates and amide and carbonyl oxygens [18,19, 25]. GB destabilizes nucleic acid duplexes at near 25 °C because its favorable interactions with nucleobase $sp^2$ C and $sp^2$ N ASA outweighs its unfavorable interaction with nucleobase $sp^2$ O ASA [18, 19, 25, 26]. GB is a protein stabilizer because its net interaction with the amide and hydrocarbon ASA exposed in unfolding is unfavorable [18, 21]. Large unfavorable interactions of GB with amide (and carboxylate) $sp^2$ O and aliphatic $sp^3$ C outweigh its favorable interactions with amide (and cationic) N and aromatic $sp^2$ C.

Classically, effects of solutes on protein stability and other processes have been quantified as thermodynamic *m*-values [27]. The *m*-value is most appropriately defined as the derivative of the observed standard free energy change ($\Delta G^o_{obs}$) for the process with respect to molal solute concentration ($m_3$) at constant temperature (T) and solution conditions [18, 28]:

$$\frac{d\Delta G^o_{obs}}{dm_3} = -RT\frac{d\ln(K_{obs})}{dm_3} = thermodynamic\ m - value \qquad \textbf{Eq. S1}$$

In Eq S1, $K_{obs}$ is the observed quotient of equilibrium product and reactant concentrations; $K_{obs}$ depends on $m_3$ because of solute effects on product vs. reactant activity coefficients. The thermodynamic *m*-value is typically independent of solute concentration over a wide range, which differs for different solutes but often extends above 1 molal.

The thermodynamic *m*-value is interpreted in terms of the interaction of the solute with the surface area that is exposed or buried in the process (i.e. the $\Delta ASA$) [18, 28]:

$$thermodynamic\ m - value = -RT\frac{d\ln(K_{obs})}{dm_3} = \sum \alpha_j \Delta ASA_j \qquad \textbf{Eq. S2}$$

In Eq. S2, $\Delta ASA_j$ is the change in water-accessible surface area of surface type j (e.g. $sp^3$ or $sp^2$ C, N or O unified atoms) and the intensity factor $\alpha_j$ is a coefficient quantifying the intrinsic strength of the interaction of the solute with a unit area of surface type *j*, determined from studies of interactions of model compounds displaying subsets of protein or nucleic acid functional groups.

A kinetic *m*-value is obtained from the derivative of an observed rate constant ($k_{obs}$) with respect to solute molality $m_3$ and interpreted analogously to a thermodynamic *m*-value.

$$kinetic\ m - value = -RT\frac{d\ln(k_{obs})}{dm_3} = \sum \alpha_j \Delta ASA_j^{\ddagger} \qquad \textbf{Eq. S3}$$

The kinetic *m*-value quantifies the effect of a solute on the height of the overall activation free energy barrier for converting reactants to the highest (rate-determining) transition state ($TS^{\ddagger}$) in the process [29, 30].

Previously-determined $\alpha_j$ - values for interactions of urea and GB with the different hybridization states of unified O, N and C atoms of proteins and nucleobases are summarized in SI Table S1. Using these $\alpha_j$ values, *m*-values quantifying effects of a solute on $K_{obs}$ or $k_{obs}$ of a protein or nucleic acid process can be interpreted to obtain information about the amount and composition of the ΔASA in that process. Conversely, *m*-values for processes can be reliably predicted if structural information is available.

For each step i of transcription initiation, the overall forward rate constant $k_{obs,i}$ is interpreted as

$$k_{obs,i} \cong K_{obs,i}^{tr} K_{obs,i}^{NTP} k_{rds,i} \qquad \textbf{Eq. S4}$$

where $K_{obs,i}^{NTP}$ and $K_{obs,i}^{tr}$ are equilibrium concentration quotients for binding the nucleotide incorporated in step i and for translocation and $k_{rds,i}$ is the rate constant for the subsequent rate-determining step (see below) [2, 3]. **Eq. S4** is applicable when both translocation and NTP binding are rapidly-established equilibria relative to the time scale of the rate-determining step and where translocation is highly disfavored ($K_{obs,i}^{tr}$<< 1) because of translocation stresses. These conditions appear applicable to initiation from the $\lambda P_R$ promoter under the conditions investigated [2, 3].

Therefore, for addition of the ith nucleotide,

$$kinetic\ m - value_i = -RT\left(\frac{\mathrm{dln}(k_{obs,i})}{dm_3}\right) \cong -RT\left(\frac{\mathrm{dln}(K_{obs,i}^{tr})}{dm_3} + \frac{dln(K_{obs,i}^{NTP})}{dm_3} + \frac{dln(k_{rds,i})}{dm_3}\right) =$$

$$\sum \alpha_j\left(\Delta ASA_{tr,j} + \Delta ASA_{NTP,j} + \Delta ASA_{rds,j}^{\ddagger}\right) \qquad \textbf{Eq. S5}$$

Goals of the urea and GB studies reported here are to test the step-wise model for disruption of contacts of RNAP with the promoter, which would allow closing of the upstream initiation bubble, and to obtain semiquantitative information about the direction and magnitude of ASA changes in these steps.

In global fitting of kinetic data at multiple solute concentrations $m_3$, the *m*-value is assumed to be constant and rate constants $k_{obs,i}^{m3}$ for each step i at different $m_3$ are constrained to be related by

$$\ln\left(k_{obs,i}^{m_3}\right) = \ln\left(k_{obs,i}^{m_{3=0}}\right) - \left(\frac{m-value}{RT}\right) m_3 \qquad \textbf{Eq. S6}$$

Values of $k_{obs,i}^{m_{3=0}}$ were obtained previously [2, 3]. In the text, we simplify the designation of observed nucleotide-incorporation 2[nd] order rate constant for step i as $k_{obs,i}$.

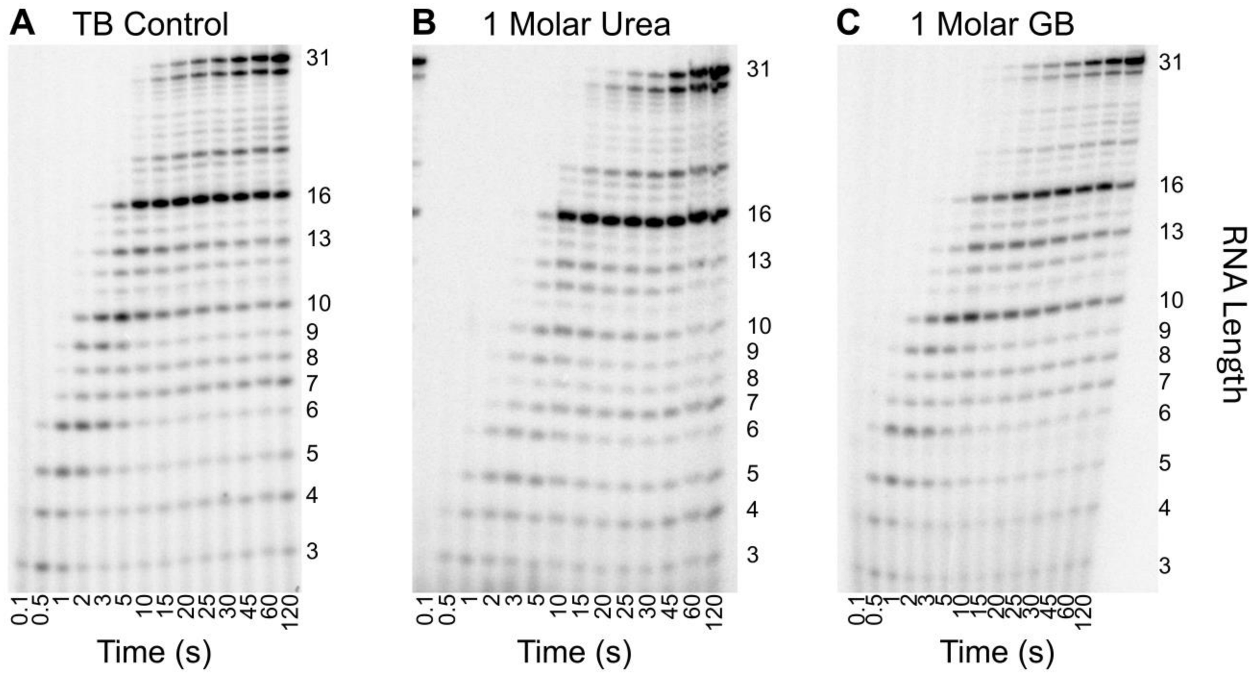

**Figure S1. Time courses of transcription initiation by E. coli RNA polymerase- $\lambda P_R$ promoter complexes at 1 M urea and 1 M GB.** Representative gel separations of short (pre-escape; <11-mer)) and long (≥11-mer) RNA transcripts as a function of time (0.1 s to 120 s) after NTP addition at the high UTP condition at 19 °C. A) Control (TB, no added solute); B) 1 M urea; C) 1 M glycine betaine (GB).

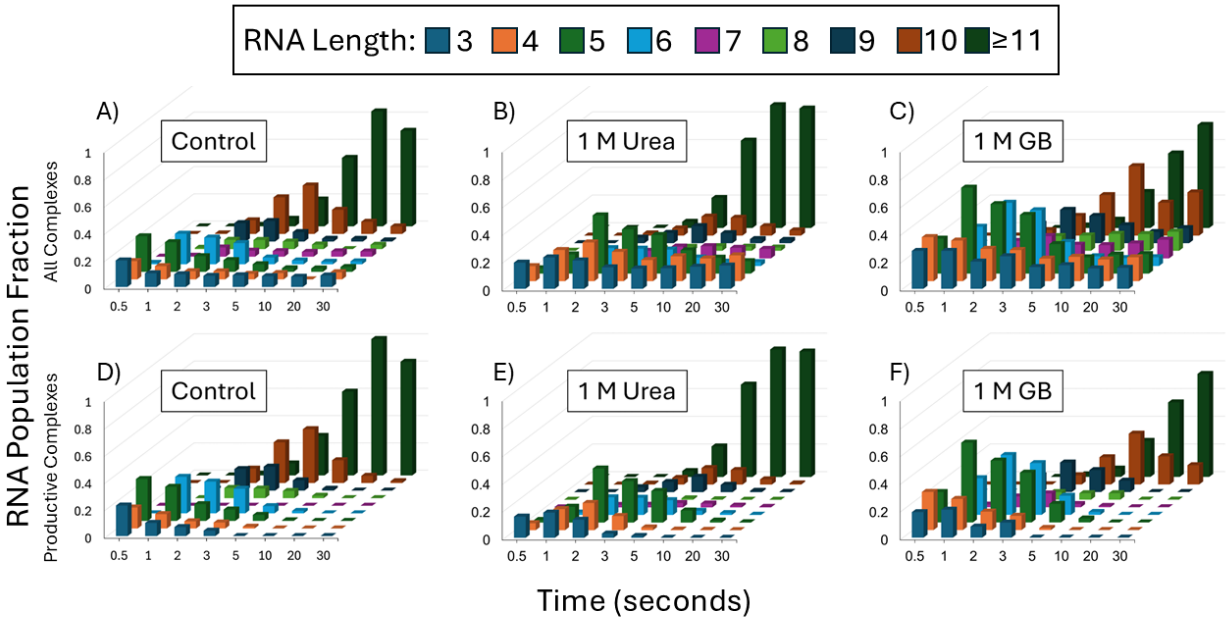

**Figure S2. Time-evolution of short RNAs (pre-escape; 3-mer to 10-mer) and of long (≥11-mer) RNA) in initiation at λP$_R$ promoter at 1.0 M Urea and 1.0 M GB.** NTP condition: high ATP, UTP, low GTP**. Panels A-C:** Total RNA from both productive and non-productive complexes. **Panels D-F**: RNA from productive complexes only. **Panels A, D:** Control (TB, no added solute [2]) **Panels B, E:** 1.0 M urea. **Panels C, F:** 1.0 M glycine betaine. For all panels, all RNA amounts are normalized to the final amount of long (≥11-mer) RNA.

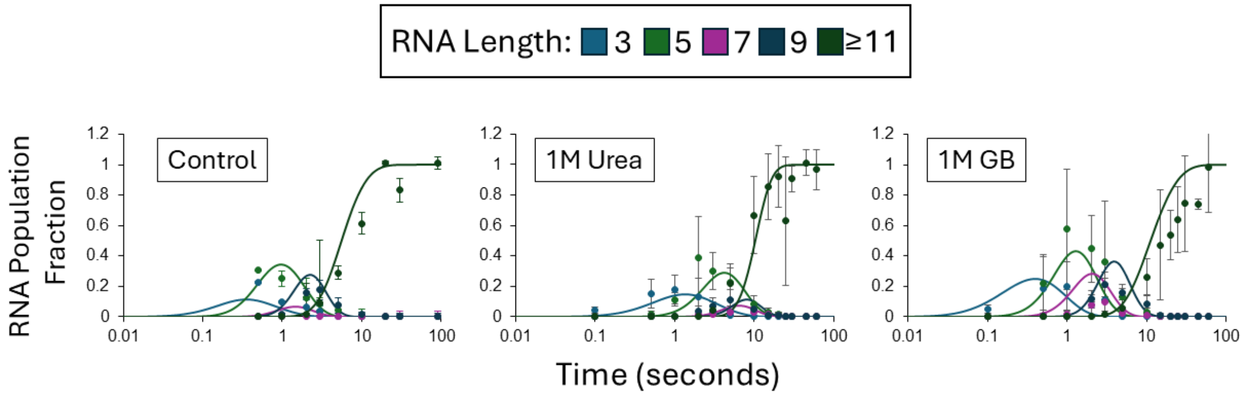

**Figure S3. Fitting of time course data to the step-by-step initiation mechanism.** Results for four pre-escape intermediates and for ≥11-mer RNA at 1.0 M urea, 1.0 M GB and the control (see **Figure S2 Panels D-F**) are compared with curves predicted from a global fit on this log time plot., **Panel A):** No added solute. **Panel B:** 1.0 M urea. **Panel C:** 1.0 M glycine betaine. All RNA amounts are normalized as in **Figure 2**.

**SI Tables**

**Table S1**: α-Values Quantifying Strengths of Interaction of Urea and Glycine Betaine (GB) With a Unit Area of Water-Accessible Surface of Different Unified Atoms of Proteins and Nucleic Acids

| Atom type | Urea α-Values<br>(cal $mol^{-1}$ $molal^{-1}$ $Å^{-2}$) | GB α-Values<br>(cal $mol^{-1}$ $molal^{-1}$ $Å^{-2}$) |
|---|---|---|
| RNAP | | |
| Amide $sp^2O$ | -0.49 ± 0.1 | 1.6 ± 0.6 |
| Carboxylate $sp^2O$ | -0.21 ± 0.09 | 1.7 ± 0.1 |
| Hydroxyl $sp^3O$ | -0.15 ± 0.03 | 0.06 ± 0.12 |
| Amide $sp^2N$ | -0.21 ± 0.13 | -1.2 ± 0.4 |
| Cationic $sp^2N$, $sp^3N$ | 0.09 ± 0.10 | -0.70 ± 0.23 |
| Aromatic $sp^2C$ | -0.52 ± 0.03 | -1.3 ± 0.2 |
| Methyl, methylene $sp^3C$ | -0.06 ± 0.03 | 0.17 ± 0.17 |
| DNA/RNA | | |
| Phosphate $sp^2O$ | -0.34 ± 0.07 | 2.8 ± 0.2 |
| Carbonyl $sp^2O$ | -0.36 ± 0.11 | 1.6 ± 0.6[c] |
| Sugar $sp^2O$, $sp^3O$ | -0.38 ± 0.04 | 1.6 ± 0.6 [c] |
| Amino $sp^3N$ | -0.16 ± 0.11 | -0.70 ± 0.23 [c] |
| Aromatic $sp^2N$ | -0.64 ± 0.04 | -1.2 ± 0.4 [c] |
| Aromatic $sp^2C$ | -0.64 ± 0.04 | -1.3 ± 0.2 [c] |
| Sugar methylene $sp^3C$ | -0.06 ± 0.03 | 0.17 ± 0.17 [c] |
| Base methyl $sp^3C$ | | |

[a] [19, 20]

[b] [19, 25]

[c] Estimated from α-values for corresponding types of protein atoms, as in [30]

**Table S2.** ASA Changes and Urea and GB *m*-Value Contributions for Disruption of Interface of $\sigma^{70}$ Regions 1.2 and 2 with Strands of Upstream Bubble (-1 to -11) DNA

| | $\Delta$ASA[a] ($\AA^2$) | Urea *m*-value contribution[b] (kcal mol$^{-1}$ molal$^{-1}$) | GB *m*-value contribution[b] (kcal mol$^{-1}$ molal$^{-1}$) |
|---|---|---|---|
| RNAP | | | |
| Aliphatic sp$^3$C | 736 | -0.05 ± 0.02 | 0.13 ± 0.13 |
| Aromatic sp$^2$C | 276 | -0.14 ± 0.01 | -0.37 ± 0.07 |
| Amide sp$^2$N | 230 | -0.05 ± 0.03 | -0.27 ± 0.1 |
| Cationic sp$^2$, sp$^3$N | 207 | 0.02 ± 0.02 | -0.14 ± 0.05 |
| Amide sp$^2$O | 82 | -0.04 ± 0.01 | 0.13 ± 0.05 |
| Carboxylate sp$^2$O | 23 | 0 ± 0 | 0.04 ± 0 |
| Hydroxyl sp$^3$O | 90 | -0.01 ± 0 | 0.01 ± 0.01 |
| Sum | 1643 | -0.28 ± 0.04 | -0.47 ± 0.19 |
| DNA | | | |
| Sugar sp$^3$C | 276 | -0.11 ± 0.01 | 0.05 ± 0.05 |
| Methyl sp$^3$C | 17 | -0.01 ± 0 | 0 ± 0 |
| Aromatic sp$^2$C | 88 | -0.06 ± 0 | -0.12 ± 0.02 |
| Aromatic sp$^2$N | 221 | -0.14 ± 0.01 | -0.26 ± 0.09 |
| Amino sp$^3$N | 168 | -0.03 ± 0.02 | -0.12 ± 0.04 |
| Carbonyl sp$^2$O | 274 | -0.1 ± 0.03 | 0.45 ± 0.16 |
| Sugar sp$^2$O | 12 | 0 ± 0 | 0.02 ± 0.01 |
| Phosphate sp$^2$O | 230 | -0.08 ± 0.02 | 0.65 ± 0.05 |
| Sum | 1285 | -0.52 ± 0.04 | 0.68 ± 0.21 |
| Total | 2929 | -0.80 ± 0.06 | 0.20 ± 0.28 |

[a] ASA calculations from 7MKD [7]

[b] Predicted from α-values in Table S1 and ΔASA information

**Table S3**. ASA Changes and *m*-Value Contributions for Disruption of Interface of $\sigma^{70}$ Regions 3 and 4 with Upstream Duplex DNA

| | $\Delta$ASA[a] ($Å^2$) | Urea *m*-value contribution[b] (kcal $mol^{-1}$ $molal^{-1}$) | GB *m*-value contribution[b] (kcal $mol^{-1}$ $molal^{-1}$) |
|---|---|---|---|
| RNAP | | | |
| Aliphatic $sp^3$C | 519 | -0.03 ± 0.02 | 0.09 ± 0.09 |
| Aromatic $sp^2$C | 34 | -0.02 ± 0 | -0.05 ± 0.01 |
| Amide $sp^2$N | 274 | -0.06 ± 0.04 | -0.32 ± 0.11 |
| Cationic $sp^2$, $sp^3$N | 144 | 0.01 ± 0.01 | -0.10 ± 0.03 |
| Amide $sp^2$O | 101 | -0.05 ± 0.01 | 0.16 ± 0.06 |
| Carboxylate $sp^2$O | 94 | -0.02 ± 0.01 | 0.16 ± 0.01 |
| Hydroxyl $sp^3$O | 57 | -0.01 ± 0 | 0 ± 0.01 |
| Sum | 1222 | -0.17 ± 0.04 | -0.05 ± 0.16 |
| DNA | | | |
| Sugar $sp^3$C | 137 | -0.05 ± 0.01 | 0.02 ± 0.02 |
| Methyl $sp^3$C | 137 | -0.1 ± 0.01 | 0.02 ± 0.02 |
| Aromatic $sp^2$C | 78 | -0.05 ± 0 | -0.10 ± 0.02 |
| Aromatic $sp^2$N | 67 | -0.04 ± 0 | -0.08 ± 0.03 |
| Amino $sp^3$N | 71 | -0.01 ± 0.01 | -0.05 ± 0.02 |
| Carbonyl $sp^2$O | 62 | -0.02 ± 0.01 | 0.10 ± 0.04 |
| Sugar $sp^2$O | 13 | 0 ± 0 | 0.02 ± 0.01 |
| Phosphate $sp^2$O | 523 | -0.18 ± 0.04 | 1.49 ± 0.12 |
| Sum | 1089 | -0.46 ± 0.04 | 1.42 ± 0.14 |
| Total | 2311 | -0.63 ± 0.06 | 1.37 ± 0.21 |

[a] ASA calculations from 7MKD [7]

[b] Predicted from α-values in Table S1 and ΔASA information

**Table S4.** ASA Changes and *m*-Value Contributions for Disruption of Interface of Core RNAP With Strands of Upstream Bubble (-1 to -11) DNA[a]

| | ΔASA[a] (Å$^2$) | Urea *m*-value contribution[b] (kcal mol$^{-1}$ molal$^{-1}$) | GB *m*-value contribution[b] (kcal mol$^{-1}$ molal$^{-1}$) |
|---|---|---|---|
| RNAP | | | |
| Aliphatic sp$^3$C | 826 | -0.05 ± 0.02 | 0.14 ± 0.15 |
| Aromatic sp$^2$C | 131 | -0.07 ± 0 | -0.17 ± 0.03 |
| Amide sp$^2$N | 488 | -0.10 ± 0.06 | -0.57 ± 0.2 |
| Cationic sp$^2$, sp$^3$N | 365 | 0.03 ± 0.04 | -0.25 ± 0.09 |
| Amide sp$^2$O | 149 | -0.07 ± 0.02 | 0.24 ± 0.09 |
| Carboxylate sp$^2$O | 88 | -0.02 ± 0.01 | 0.15 ± 0.01 |
| Hydroxyl sp$^3$O | 73 | -0.01 ± 0 | 0 ± 0.01 |
| Sum | 2120 | -0.29 ± 0.08 | -0.46 ± 0.28 |
| DNA | | | |
| Sugar sp$^3$C | 328 | -0.13 ± 0.01 | 0.06 ± 0.06 |
| Methyl sp$^3$C | 94 | -0.07 ± 0.01 | 0.02 ± 0.02 |
| Aromatic sp$^2$C | 148 | -0.09 ± 0.01 | -0.2 ± 0.04 |
| Aromatic sp$^2$N | 183 | -0.12 ± 0.01 | -0.21 ± 0.08 |
| Amino sp$^3$N | 170 | -0.03 ± 0.02 | -0.12 ± 0.04 |
| Carbonyl sp$^2$O | 340 | -0.12 ± 0.04 | 0.55 ± 0.2 |
| Sugar sp$^2$O | 31 | -0.01 ± 0 | 0.05 ± 0.02 |
| Phosphate sp$^2$O | 545 | -0.18 ± 0.04 | 1.55 ± 0.13 |
| Sum | 1512 | -0.62 ± 0.06 | 1.64 ± 0.26 |
| Total | 3632 | -0.92 ± 0.10 | 1.18 ± 0.38 |

[a] ASA calculations from 7MKD [7]

[b] Predicted from α-values in Table S1 and ΔASA information

**Table S5.** ASA Changes and *m*-Value Contributions for Disruption of Interface of $\sigma^{70}$ Subunit with Core RNAP

| | $\Delta$ASA[a] ($Å^2$) | Urea *m*-value contribution[b] (kcal $mol^{-1}$ $molal^{-1}$) | GB *m*-value contribution[b] (kcal $mol^{-1}$ $molal^{-1}$) |
|---|---|---|---|
| Aliphatic $sp^3C$ | 5759 | -0.37 ± 0.17 | 1.0 ± 1.0 |
| Aromatic $sp^2C$ | 477 | -0.25 ± 0.01 | -0.64 ± 0.11 |
| Amide $sp^2N$ | 1118 | -0.24 ± 0.15 | -1.3 ± 0.46 |
| Cationic $sp^2$, $sp^3N$ | 544 | 0.05 ± 0.05 | -0.38 ± 0.13 |
| Amide $sp^2O$ | 1007 | -0.50 ± 0.11 | 1.64 ± 0.60 |
| Carboxylate $sp^2O$ | 865 | -0.19 ± 0.08 | 1.46 ± 0.10 |
| Hydroxyl $sp^3O$ | 439 | -0.06 ± 0.02 | 0.03 ± 0.05 |
| Sum | 10208 | -1.55 ± 0.27 | 1.81 ± 1.29 |

[a] ASA calculations from 7MKD [7]

[b] Predicted from α-values in Table S1 and ΔASA information

**Table S6**. ASA Changes and m-Value Contributions for Binding the First Initiating NTP (ATP)

| RNAP | | | |
|---|---|---|---|
| Surface | - ΔASA (Å$^2$)[a] | Urea *m*-value[b] contribution (kcal mol$^{-1}$ molal$^{-1}$) | GB *m*-value[b] contribution (kcal mol$^{-1}$ molal$^{-1}$) |
| Carboxylate sp$^2$O | 28 | 0.01 ± 0 | -0.05 ± 0 |
| Amide sp$^2$O | 27 | 0.01 ± 0 | -0.04 ± 0.02 |
| Amide sp$^2$N | 57 | 0.01 ± 0.01 | 0.07 ± 0.02 |
| Cationic sp$^2$ N, sp$^3$N | 44 | 0 ± 0 | 0.03 ± 0.01 |
| Methyl, methylene sp$^3$ C | 41 | 0 ± 0 | -0.01 ± 0.01 |
| Aromatic sp$^2$C | 20 | 0.01 ± 0 | 0.03 ± 0 |
| Total RNAP | 217 Å$^2$ | 0.04 ± 0.01 | 0.03 ± 0.03 |
| ATP | | | |
| Surface | - ΔASA (Å$^2$)[a] | Urea *m*-value contribution (kcal mol$^{-1}$ molal$^{-1}$) | GB *m*-value contribution (kcal mol$^{-1}$ molal$^{-1}$) |
| Phosphate sp$^2$O | 130 | 0.04 ± 0.01 | -0.37 ± 0.03 |
| Carbonyl sp$^2$O | 31 | 0.01 ± 0 | -0.05 ± 0.02 |
| Sugar sp$^2$O, sp$^3$O | 10 | 0 ± 0 | -0.02 ± 0.01 |
| Amino sp$^3$N | 52 | 0.01 ± 0.01 | 0.04 ± 0.01 |
| Aromatic sp$^2$N | 50 | 0.03 ± 0 | 0.06 ± 0.02 |
| Aromatic sp$^2$C | 85 | 0.05 ± 0 | 0.11 ± 0.02 |
| Sugar sp$^3$C | 43 | 0.02 ± 0 | -0.01 ± 0.01 |
| Total ATP | 402 Å$^2$ | 0.17 ± 0.01 | -0.24 ± 0.05 |
| Total RNAP + ATP | 619 Å$^2$ | 0.21 ± 0.02 | -0.21 ± 0.06 |

[a] ASA calculations from 4Q4Z [8]

[b] Predicted from α-values in Table S1 and ΔASA information

**Table S7.** ASA Changes and *m*-Value Contributions for Binding the Second Initiating NTP (CTP analog CMPCPP)

| RNAP | | | |
|---|---|---|---|
| Surface | ΔASA (Å$^2$)[a] | Urea *m*-value contribution[b] (kcal mol$^{-1}$ molal$^{-1}$) | GB m-value contribution[b] (kcal mol$^{-1}$ molal$^{-1}$) |
| Amide sp$^2$O | 170 | 0.08 ± 0.02 | -0.28 ± 0.10 |
| Carboxylate sp$^2$O | 103 | 0.02 ± 0.01 | -0.17 ± 0.01 |
| Hydroxyl sp$^3$O | 16 | 0 ± 0 | 0 ± 0 |
| Amide sp$^2$N | 114 | 0.02 ± 0.01 | 0.13 ± 0.05 |
| Cationic sp$^2$N, sp$^3$N | 35 | 0 ± 0 | 0.02 ± 0.01 |
| Methyl, methylene sp$^3$C | 441 | 0.03 ± 0.01 | -0.08 ± 0.08 |
| Aromatic sp$^2$C | -28 | -0.01 ± 0 | -0.04 ± 0.01 |
| TOTAL RNAP | 851 Å$^2$ | 0.14 ± 0.03 | -0.41 ± 0.14 |
| Nucleotide (CMPCPP, ATP)[c] | | | |
| Surface | ΔASA (Å$^2$)[a] | Urea *m*-value contribution[b] (kcal mol$^{-1}$ molal$^{-1}$) | GB *m*-value contribution[b] (kcal mol$^{-1}$ molal$^{-1}$) |
| Phosphate sp$^2$O | 201 | 0.07 ± 0.01 | -0.57 ± 0.05 |
| Carbonyl sp$^2$O | 50 | 0.02 ± 0.01 | -0.08 ± 0.03 |
| Sugar sp$^2$O, sp$^3$O | 1 | 0 ± 0 | 0 ± 0 |
| Amino sp$^3$N | 59 | 0.01 ± 0.01 | 0.04 ± 0.01 |
| Aromatic sp$^2$N | 25 | 0.02 ± 0 | 0.03 ± 0.01 |
| Aromatic sp$^2$C | 93 | 0.06 ± 0 | 0.12 ± 0.02 |
| Sugar methylene sp$^3$C | 65 | 0.03 ± 0 | -0.01 ± 0.01 |
| Total CMPCPP | 494 Å$^2$ | 0.20 ± 0.02 | -0.47 ± 0.06 |
| Total RNAP + CMPCPP | 1345 Å$^2$ | 0.34 ± 0.03 | -0.88 ± 0.15 |

[a] ASA calculations from 4Q4Z [8]

[b] Predicted from α-values in Table S1 and ΔASA information

[c] Nucleotide contributions include changes in ATP ASA in binding CMPCPP.

**Table S8.** ASA Changes and m-Value Contributions for Converting the Partially Folded Trigger Loop to Fully-Folded Trigger Helices

| RNAP | | | |
|---|---|---|---|
| Surface | -ΔASA (Å$^2$)[a] | Urea *m*-value contribution[b] (kcal mol$^{-1}$ molal$^{-1}$) | GB *m*-value contribution[b] (kcal mol$^{-1}$ molal$^{-1}$) |
| Amide sp$^2$O | 62 | 0.03 ± 0.01 | -0.10 ± 0.04 |
| Hydroxyl sp$^3$O | 18 | 0 ± 0 | 0 ± 0 |
| Carboxylate sp$^2$O | -27 | -0.01 ± 0 | 0.05 ± 0 |
| Amide sp$^2$N | 63 | 0.01 ± 0.01 | 0.08 ± 0.03 |
| Cationic sp$^2$N, sp$^3$N | 3 | 0 ± 0 | 0 ± 0 |
| Methyl, methylene sp$^3$C | 74 | 0.01 ± 0 | -0.01 ± 0.01 |
| Aromatic sp$^2$C | 53 | 0.03 ± 0 | 0.07 ± 0.01 |
| Total RNAP | 246 Å$^2$ | 0.07 ± 0.01 | 0.09 ± 0.05 |
| Nucleotide (CMPCPP, ATP) | | | |
| Surface | -ΔASA (Å$^2$)[a] | Urea *m*-value contribution[b] (kcal mol$^{-1}$ molal$^{-1}$) | GB *m*-value contribution[b] (kcal mol$^{-1}$ molal$^{-1}$) |
| Phosphate sp$^2$O | 79 | 0.03 ± 0.01 | -0.22 ± 0.02 |
| Sugar sp$^2$O, sp$^3$O | 3 | 0 ± 0 | 0 ± 0 |
| Carbonyl sp$^2$O | 0 | 0 ± 0 | 0 ± 0 |
| Amino sp$^3$N | 4 | 0 ± 0 | 0 ± 0 |
| Aromatic sp$^2$N | 0 | 0 ± 0 | 0 ± 0 |
| Aromatic sp$^2$C | 11 | 0.01 ± 0 | 0.01 ± 0 |
| Sugar methylene sp$^3$C | -4 | 0 ± 0 | 0 ± 0 |
| Total CMPCPP | 93 Å$^2$ | 0.03 ± 0.01 | -0.21 ± 0.02 |
| Total RNAP + CMPCPP | 339 Å$^2$ | 0.10 ± 0.01 | -0.13 ± 0.05 |

[a] ASA calculations from 4Q4Z [8]

[b] Predicted from α-values in Table S1 and ΔASA information

**Table S9.** Predicted ASA Changes and *m*-Value Contributions for Overall Step of 2-mer (pppApU) Synthesis

| RNAP | | | |
|---|---|---|---|
| Surface | -ΔASA (Å$^2$)[a] | Urea *m*-value contribution[b] (kcal mol$^{-1}$ molal$^{-1}$) | GB *m*-value contribution[b] (kcal mol$^{-1}$ molal$^{-1}$) |
| Amide sp$^2$O | 231 | 0.11 ± 0.02 | -0. 37 ± 0.11 |
| Carboxylate sp$^2$O | 76 | 0.02 ± 0.01 | -0.13 ± 0.01 |
| Hydroxyl sp$^3$O | 34 | 0.01 ± 0 | 0 ± 0 |
| Amide sp$^2$N | 177 | 0.04 ± 0.02 | 0.21 ± 0.05 |
| Cationic sp$^2$N, sp$^3$N | 38 | 0 ± 0 | 0.03 ± 0.01 |
| Methyl, methylene sp$^3$ C | 515 | 0.03 ± 0.01 | -0.09 ± 0.08 |
| Aromatic sp$^2$C | 24 | 0.01 ± 0 | 0.03 ± 0.01 |
| Total RNAP | 1095 | 0.22 ± 0.03 | -0.32 ± 0.15 |
| Nucleotide (CMPCPP, ATP) | | | |
| Surface | -ΔASA (Å$^2$)[a] | Urea *m*-value contribution[b] (kcal mol$^{-1}$ molal$^{-1}$) | GB *m*-value contribution[b] (kcal mol$^{-1}$ molal$^{-1}$) |
| Phosphate sp$^2$O | 280 | 0.10 ± 0.02 | -0.78 ± 0.05 |
| Carbonyl sp$^2$O | 50 | 0.02 ± 0.01 | -0.08 ± 0.03 |
| Sugar sp$^2$O, sp$^3$O | 4 | 0 ± 0 | 0.01 ± 0.01 |
| Amino sp$^3$N | 63 | 0.01 ± 0.01 | 0.04 ± 0.01 |
| Aromatic sp$^2$N | 25 | 0.02 ± 0 | 0.03 ± 0.01 |
| Aromatic sp$^2$C | 104 | 0.07 ± 0 | 0.14 ± 0.02 |
| Sugar methylene sp$^3$C | 61 | 0.02 ± 0 | -0.01 ± 0.01 |
| Total CMPCPP | 587 | 0.24 ± 0.02 | -0.65 ± 0.07 |
| Total RNAP + CMPCPP | 1682 | 0.46 ± 0.04 | -0.97 ± 0.16 |

[a] ASA calculations from 4Q4Z [8]

[b] Predicted from $\alpha$-values in Table SX and ΔASA information

[c] Nucleotide contributions include changes in ATP ASA in binding CMPCPP.

**Table S10.** Predicted Contributions to Urea and GB m-Values for Different Models for Timing of Bubble Collapse and/or Sigma Release

| Process | $\Delta$ASA (Å$^2$) | Urea *m*-value (kcal mol$^{-1}$ molal$^{-1}$) | GB *m*-value (kcal mol$^{-1}$ molal$^{-1}$) |
|---|---|---|---|
| A) Disrupting $\sigma^{70}$ Contacts with ss DNA (-1 to -11) | 2.9 x 10$^3$ | – 0.8 ± 0.1 | + 0.2 ± 0.3 |
| B) Disrupting Core RNAP Contacts with ss DNA | 3.6 x 10$^3$ | – 0.9 ± 0.1 | + 1.2 ± 0.4 |
| C) 12-mer Duplex Formation from Half-Stacked Strands | -- | + 1.2 ± 0.4 | + 0.3 ± 0.1 |
| D) Disrupting $\sigma^{70}$ Contacts with ds DNA (-12 to -38?) | 2.3 x 10$^3$ | – 0.6 ± 0.1 | + 1.4 ± 0.2 |
| E) Disrupting $\sigma^{70}$ Contacts with Core RNAP | 1.0 x 10$^4$ | -1.6 ± 0.3 | + 1.8 ± 1.3 |
| Sum of A, B, C, D, E above | | - 2.7 ± 0.5[a] | + 4.9 ± 1.7[b] |
| Sum of Excess m-Values for steps 5 - 11 | | - 1.9 ± 0.8 | + 4.4 ± 0.5 |
| Without C | | - 3.1 ± 0.3[a] | + 4.6 ± 1.7[b] |
| Without E | | - 1.1 ± 0.4[c] | + 3.1 ± 0.5[d] |
| Without C and E | | - 2.3 ± 0.2[a] | + 2.8 ± 1.7[d] |
| Sum of Excess m-Values for steps 10 - 11 | | -2.3 ± 0.2 | + 1.2 ± 0.1 |
| Sum of D, E above | | - 2.2 ± 0.3[a] | + 3.2 ± 1.3[d] |
| Sum of C, D, E above | | - 1.0 ± 0.5[c] | + 3.5 ± 1.7[d] |
| D only | | - 0.6 ± 0.1[c] | + 1.4 ± 0.2[b] |
| Sum of C, D above | | + 0.6 ± 0.4[c] | + 1.7 ± 0.2[b] |
| Sum of Excess m-values for Steps 5 - 9 | | + 0.4 ± 0.8 | + 3.2 ± 0.4 |
| Sum of A, B, C above | | - 0.5 ± 0.4[a] | + 1.7 ± 0.5[d] |
| Sum of A, B above | | - 1.7 ± 0.1[c] | + 1.4 ± 0.5[d] |

[a] Predicted urea kinetic *m*-value agrees with observed within the combined uncertainties
[b] Predicted GB kinetic *m*-value agrees with observed within the combined uncertainties
[c] Predicted urea kinetic *m*-value disagrees with observed by more than the combined uncertainties
[d] Predicted GB kinetic *m*-value disagrees with observed by more than the combined uncertainties

**Table S11.** Predicted Contributions to Excess Activation Energies for Different Models for Timing of Bubble Collapse

| Excess $E_{act}$ (kcal $mol^{-1}$) | Steps 5 - 11 | Steps 10 - 11 | Steps 5 - 9 |
|---|---|---|---|
| Observed | - 124 ± 18 | - 15 ± 5 | -109 ± 17 |
| Predicted for duplex formation in steps 5 - 11 | - 120 ± 20[a] | NA[b] | NA[b] |
| After step 11 | ~ 0 | ~ 0 | ~ 0 |
| In steps 10 -11 only | - 120 ± 20 | -120 ± 20 | ~ 0 |
| In steps 5 – 9 only | - 120 ± 20 | ~ 0 | - 120 ± 20 |

[a] Assuming bases in open strands are half-stacked [3]

[b] NA, not applicable
